# The genome sequence of the Violet Carpenter Bee, *Xylocopa violacea* (Linnaeus, 1785): a hymenopteran species undergoing range expansion

**DOI:** 10.1101/2024.04.03.587942

**Authors:** Will J Nash, Angela Man, Seanna McTaggart, Kendall Baker, Tom Barker, Leah Catchpole, Alex Durrant, Karim Gharbi, Naomi Irish, Gemy Kaithakottil, Debby Ku, Aaliyah Providence, Felix Shaw, David Swarbreck, Chris Watkins, Ann M. McCartney, Giulio Formenti, Alice Mouton, Noel Vella, Björn M von Reumont, Adriana Vella, Wilfried Haerty

**Affiliations:** The Earlham Institute, Norwich Research Park, Colney Lane, Norwich, NR4 7UZ, UK; Genomics Institute, University of California, Santa Cruz, CA 95060, USA; The Vertebrate Genome Laboratory, The Rockefeller University, 1240 York Ave, 10065 New York, USA; Department of Biology, University of Florence, Sesto Fiorentino, Italy; InBios - Conservation Genetics Laboratory, University of Liege, Chemin de la Vallee 4, 4000 Liege, Belgium; SEED - Departement des sciences et gestion de l’environnement, University of Liege, Chemin de la Vallee 4, 4000 Liege, Belgium; Conservation Biology Research Group, Biology Department, University of Malta, Msida, MSD 2080, Malta; LOEWE Center for Translational Biodiversity Genomics (LOEWE-TBG), Senckenberganlage 25, 60325 Frankfurt, Germany; Applied Bioinformatics Group, Faculty of Biological Sciences, Goethe University Frankfurt, Max-von-Laue-Str. 13, 60438 Frankfurt, Germany; School of Biological Sciences, The University of East Anglia, Norwich, NR4 7TJ, UK

**Keywords:** Hymenoptera, Genome, Long-read, Assembly, Hi-C, Repetitive DNA

## Abstract

We present a reference genome assembly from an individual male Violet Carpenter Bee (*Xylocopa violacea,* Linnaeus, 1758). The assembly is 1.02 gigabases in span. 48% of the assembly is scaffolded into 17 pseudo-chromosomal units. The mitochondrial genome has also been assembled and is 21.8 kilobases in length. The genome is highly repetitive, likely representing a highly heterochromatic architecture expected of bees from the genus *Xylocopa*. We also use an evidence-based methodology to annotate 10,152 high confidence coding genes. This genome was sequenced as part of the pilot project of the European Reference Genome Atlas (ERGA) and represents an important addition to the genomic resources available for Hymenoptera.

## Introduction

We live in a time of unprecedented biodiversity loss (Ceballos and Ehrlich, 2023) exemplified by the global decline of insect fauna undeniably associated with anthropogenic stressors (Outhwaite *et al*., 2022). Insect biodiversity loss puts key ecosystem services, such as pollination (Ollerton, 2021) and decomposition (Yang and Gratton, 2014), at risk. Although there is strong evidence of insect declines in the recent history (Hallmann *et al*., 2017; Powney *et al*., 2019), changes in global climate have also seen patterns of range shift in many taxa (e.g. Kerr *et al*., 2015; Lehmann *et al*., 2020; Rollin *et al*., 2020; Halsch *et al*., 2021; Skendžić *et al*., 2021). The European Reference Genome Atlas (ERGA, Mc Cartney *et al*., 2023) aims to empower research communities to expand the taxonomic coverage of genomic resources, enabling cross taxa analyses to address continent-scale questions, such as those surrounding range shifts, at the genomic level.

There are currently no annotated, reference quality, genomic resources for the Carpenter bees (Hymenoptera: Apidae). They are classified as a single genus, *Xylocopa* (Latreille, 1802), which contains around 400 species (Gerling *et al*., 1989; Leys *et al*., 2000, 2002; Michener, 2007), and are considered as essential pollinators (e.g. Vargas *et al*., 2017; Malabusini *et al*., 2019). In Europe, the most widespread *Xylocopa* species is the Violet Carpenter Bee, *Xylocopa violacea* (Linnaeus, 1758) (Vicidomini, 1996). This species has a pan-European distribution (Figure 1, https://www.gbif.org/species/1342108) that also extends to Algeria and Turkey (Gerling *et al*., 1989; Aouar-Sadli *et al*., 2008; Tezcan and Skyrpan, 2022), Iraq and India (Dar *et al*., 2016; Bamarni and Elsaiegh, 2022).

**Figure 1.**
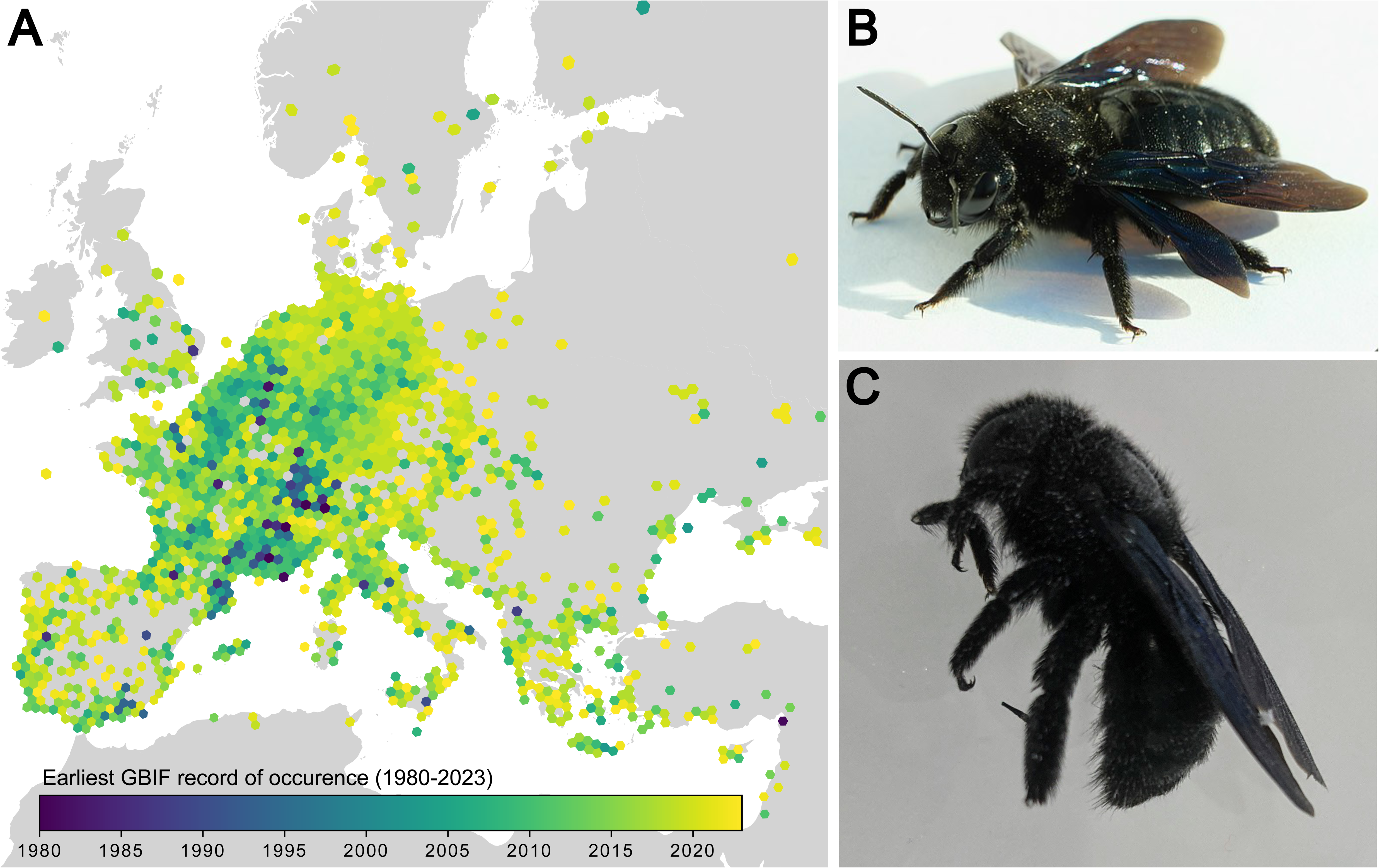
The Violet Carpenter Bee, *Xylocopa violacea*. **A)** Records of *X. violacea* occurrence in Europe between 1980 and 2023 (GBIF.org, 04 December 2023, https://doi.org/10.15468/dl.3gr8wv). Hexes are coloured by earliest year of occurrence; lighter colours are more recent. Records prior to 1980 not plotted. **B)** A female *X. violacea* individual (Bautsch, CC0, via Wikimedia Commons.) **C)** The male *X. violacea* (iyXylViol4) used for DNA sequencing in this study.

In recent years, *Xylocopa violacea* has exhibited a marked range expansion, with records in Germany (Praz *et al*., 2022), Czech Republic (Kleprlíková and Vrabec, 2020), Poland (Banaszak *et al*., 2019), and as far north as Sweden (Cederberg and Others, 2018) (Figure 1). The northward expansion of the Violet Carpenter Bee’s range may be attributed to various factors, including climatic changes in Europe (Banaszak *et al*., 2019). *Xylocopa violacea* is a solitary bee (Vicidomini, 1996), although within the genus there is evidence for several independent transitions to sociality (Gerling *et al*., 1989; Sless and Rehan, 2023). *X. violacea* also exhibits a lineage specific microbiome (Alberoni *et al*., 2019; Holley *et al*., 2022; Handy *et al*., 2023) and a distinctive venom profile with novel melittin variants that show potential for anticancer applications (von Reumont *et al*., 2022; Erkoc *et al*., 2022). There is only a contig-level assembly of the *X. violacea* genome currently available (Koludarov *et al*., 2023).

Here, we present a pseudo-chromosomal assembly of the genome of *Xylocopa violacea.* The genome was sequenced as part of the pilot project of the ERGA (Mc Cartney *et al*., 2023). The ERGA consortium is pioneering a democratised approach to biodiversity sequencing, and paired a sample ambassador from Malta, where *X. violacea* is an important and understudied species, with a sequencing centre in the UK order to generate the assembly presented here. The *X. violacea* genome assembly is characterised by its highly heterochromatic karyotype, a trait also shared by other *Xylocopa* species (Hoshiba and Imai, 1993). This genomic resource fills an important gap in the taxonomy of the Apidae, and also releases the potential to study the expanding population of this important pollinating species at the genomic level (e.g. Formenti *et al*., 2022; Webster *et al*., 2022).

## Materials and Methods

### Sample Acquisition

A male (iyXylViol4, ERS10526494) and female (iyXylViol2, ERS10526492) *Xylocopa violacea* individual were collected at Chadwick Lakes, Rabat, Malta (Latitude: 35.894639, Longitude: 14.392165). Samples were chilled to 4°C, preserved in dry ice, and maintained at −80°C until shipment to the Earlham Institute, Norwich, UK following Nagoya Protocol, permit ABSCH-IRCC-MT-255778-1. Sample metadata conformed to ERGA sample manifest standards (Böhne *et al*., 2024) and were submitted to ENA using COPO (Shaw *et al*., 2020).

### DNA Library Preparation and Sequencing

High molecular weight (HMW) DNA was extracted from thorax tissue of an individual male bee (iyXylViol4) using the Qiagen MagAttract HMW DNA Kit, with modifications as described in Mullin *et al*. (2022). HiFi library preparation and Pacific Biosciences (PacBio) sequencing were carried out following the low-input protocol described in Mullin *et al*. (2022), (Supplementary Methods) and sequenced on four Sequel II SMRT^®^ Cell 8M (diffusion loading, 30-hour movie, 2-hour immobilisation time, 2-hour pre-extension time, 60-77 pM on plate loading concentration).

### RNA Extraction, RNA-seq Library Preparation and Sequencing

RNA extractions were conducted on flash frozen head, thorax, abdomen, and leg tissues from an individual female bee (iyXylViol2) using the Omega EZNA Total RNA Kit I (R6834-01). RNA-seq libraries were then constructed using the NEBNext Ultra II RNA Library prep for Illumina kit (NEB#E7760L) NEBNext Poly(A) mRNA Magnetic Isolation Module (NEB#7490) and NEBNext Multiplex Oligos for Illumina^®^ (96 Unique Dual Index Primer Pairs) (E6440S) at a concentration of 10uM. Libraries were sequenced on an SP flow cell on a NovaSeq 6000 instrument set up to sequence 150bp paired end reads.

### Iso-Seq Library Preparation and Sequencing

PacBio Iso-Seq libraries were constructed starting from 234-300 ng of total RNA from the 4 tissue specific extractions described above. Reverse transcription cDNA synthesis was performed using NEBNext® Single Cell/Low Input cDNA Synthesis & Amplification Module (NEB, E6421). Samples were barcoded and the library pool was prepared according to the guidelines laid out in the Iso-Seq protocol version 02 (PacBio, 101-763-800), using SMRTbell express template prep kit 2.0 (PacBio, 102-088-900). The Iso-Seq pool was sequenced on the PacBio Sequel II instrument with one Sequel II SMRT^®^ Cell 8M.

### Hi-C Library Preparation and Sequencing

High-throughput/resolution chromosome conformation capture-based (Hi-C) sequencing data was generated from head tissue of male individual iyXylViol4 using the Arima Genome Wide Hi-C kit, the NEBNext Ultra II DNA Library preparation kit, and Kappa HiFi HotStart ReadyMix. The resulting libraries were sequenced on an SP flow cell, on the Novaseq 6000 instrument, sequencing 150bp paired end reads.

### Contig level genome assembly

HiFi reads were extracted from the raw Pacific Biosciences output by the Earlham Institute core bioinformatics group using the Pacific Biosciences SMRTlink pipeline (v10.1.0.119588). Prior to assembly, HiFi reads were trimmed for adapter sequences with Cutadapt (v3.2, Martin, 2011). The genome was assembled with hifiasm (v0.18.5, Cheng *et al*., 2021). Mitochondrial contigs were identified with MitoHifi (v3.0.0, Uliano-Silva *et al*., 2023), using the *Apis mellifera* mitochondrial genome (<u>OK075087.1</u>) as a closely related guide. All putative mitochondrial contigs were removed prior to scaffolding, and the MitoHifi best fit mitochondrial sequence was added back into the assembly following scaffolding. Contaminant contigs were identified and removed as the intersect of the outputs of Kraken2 (v2.0.7, Wood *et al*., 2019), BlobTools (v1.1.1, Laetsch and Blaxter, 2017), barnapp (v0.9, Table S1), CAT (v5.2.3,von Meijenfeldt *et al*., 2019), and FCS-GX (v0.3.0, Astashyn *et al*., 2023). Assembly completeness was assessed with BUSCO (v5.0.0, Manni *et al*., 2021) using hymenoptera_odb10. Assembly quality and kmer completeness were assessed with Merqury (v1.3, Rhie *et al*., 2020). Genome size of the final assembly was estimated using FastK (Table S1) and GeneScopeFK (Table S1).

### Hi-C Read QC & Scaffolding

Raw Hi-C reads were trimmed for adapters using trimmomatic (v0.39, Bolger *et al*., 2014) with the adapters.fa file from bbmap (v35.85, Bushnell, 2014) as input (see Supp. Methods). Hi-C reads were mapped to the draft assembly with Juicer (v1.6, Durand *et al*., 2016). Following the removal of contigs assigned as contaminant or mitochondrial, Hi-C reads were mapped to the resulting assembly using the Arima Mapping Pipeline (Table S1). The resulting mappings were used to scaffold the decontaminated assembly using YaHS (v1.2a.2, Zhou *et al*., 2023).

### Manual Curation of Scaffolded Assembly

Following scaffolding, trimmed, unfiltered Hi-C reads were mapped to the scaffolded assembly using Juicer (v1.6, Durand *et al*., 2016). Using these mappings, the scaffolded assembly was manually curated to pseudo-chromosomal level using Pretext-Map (v0.1.9, Table S1) contact maps visualised in PretextView (v0.2.5, Table S1). Inputs for PrextextView (Coverage track, Gap track, Telomere track) were created using the eihic pipeline (Table S1) in curation mode (-*c*). Following curation, the Rapid Curation Pipeline (Table S1), developed by the GRiT team at the Wellcome Sanger Institute, was used to extract the manually curated assembly in fasta format.

### Annotation

Annotation of repetitive DNA content was performed using the EI-Repeat pipeline (v1.3.4, Table S1) which uses third party tools for repeat calling. The repeat content of the iyXylViol4 assembly was further classified using srf (Zhang *et al*., 2023) and TRASH (Wlodzimierz *et al*., 2023), <u>and visualised using StainedGlass</u> (Vollger *et al*., 2022). The telomeric repeat landscape was explored using the explore and search functions of tidk (Table S1). Gene models were generated from the iyXylViol4 assembly using REAT - Robust and Extendable eukaryotic Annotation Toolkit (Table S1) and Minos (Table S1) which mayke use of Mikado (Table S1), Portcullis (Table S1) and many third-party tools (listed in the above repositories).

## Results & Discussion

### DNA sequencing

HMW DNA extractions from two 30 mg sections of thorax tissue from a single male *Xylocopa violacea* individual (iyXylViol4) yielded 829 ng of HMW DNA, with 74-84% of fragments over 40 kb fragment size (Figure S1). Following library preparation, 2,520,442 PacBio HiFi Reads were obtained (21.8x coverage of the final assembly). The whole head tissue from this individual (98mg) was used to generate 535,271,589 Illumina short reads following proximity ligation and Arima High Coverage Hi-C library preparation (see Supp. Results). Sequencing of this library produced 509,760,108 read pairs.

### Transcriptome sequencing

Total RNA was extracted from four tissues segments (Head, Thorax, Abdomen, Legs) from a second individual (female, iyXylViol2). These tissues produced 4.3 µg, 3.6 µg, 18.2 µg, 2.6 µg of total RNA respectively. We generated 149,032,417, 107,159,638, 116,609,061, and 148,189,077 Illumina RNA-seq short reads respectively for the head, thorax, abdomen, and legs. Additional RNA-seq reads, from *X. violacea* venom gland, were downloaded from SRA (SRR14690757, Koludarov *et al*., 2023). The same extractions were also used to generate 790,150; 717,956; 977,170, and 999,264 PacBio Iso-Seq long reads for the head, thorax, abdomen, and legs respectively. Cumulatively, this represented an average of 81.76x long-read coverage of the transcriptome.

### Genome Assembly

The initial contig assembly had 1224 contigs and spanned 1.08 Gb with an N50 of 5.91 Mb (Table 1). Prior to scaffolding, 161 contigs (59.8 Mb) were classified as contaminant content and removed from the assembly. A contig was only classified as contaminant and removed if it was identified in the output of 2 of the following tools: Contigs identified as not within the Instecta by Kraken2 (316), contigs classified as ‘’no-hit’ by blobtools (389), contigs identified as bacterial or archaeal 16s by barnapp (384), contigs classified as bacterial or viral by CAT (4), or contigs identified as contaminants by FCS-GX (1). For further details see Table S6. 79 mitochondrial candidates (1.7 Mb), identified by MitoHifi, were also removed. With this content removed, the assembly had 984 contigs spanning 1.02 Gb, with an N50 of 5.96 Mb (Table 1).

**Table 1.**
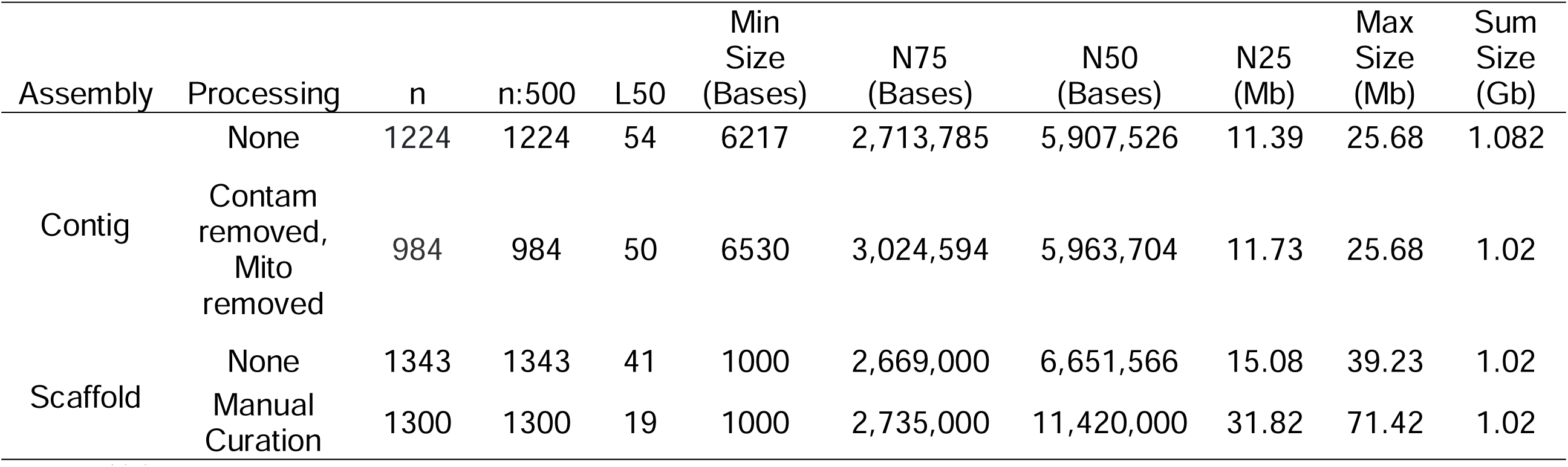
Contiguity statistics of the iyXylViol4 assembly at four stages of the assembly pipeline. Statistics generated using abyss-fac (Jackman *et al*., 2017). Contam = Contigs identified as contaminant, see main text, Mito = putuatuive mitochondrial contigs, identified using MitoHifi (Uliano-Silva *et al*., 2023), see main text.

Scaffolding generated an assembly with 1343 scaffolds spanning 1.02 Gb with an N50 of 6.65 Mb (Table 1). The scaffolded assembly was manually curated to give the final pseudo-chromosomal iyXylViol4 assembly (<u>GCA_963969225.1</u>), containing 1300 scaffolds over 1.02 Gb, and an N50 of 11.42 Mb (Figure 2, Table 1). The consensus mitogenome (21.8 Kb) was added to the assembly following manual curation and annotation. The iyXylViol4 assembly contains 17 pseudo-chromosomal units. One of these units has Hi-C telomeric signal at both ends, and the remaining 16 of which have Hi-C telomeric signal at one end. *Xylocopa violacea* has been suggested to have a karyotype of 16 (Granata, 1909), similar to a related species, *X. fenestra*, (Kumbkarni, 1965; Kerr and da Silveira, 1972), thus it is possible that two of the remaining super scaffolds in the iyXylViol4 assembly correspond to chromosomal arms with insufficient Hi-C signal to be joined. Alternatively, *X. appendiculata* has a karyotype of 17 chromosomes including a majority of pseudo-acrocentric chromosomal morphologies (Hoshiba and Imai, 1993).

**Figure 2.**
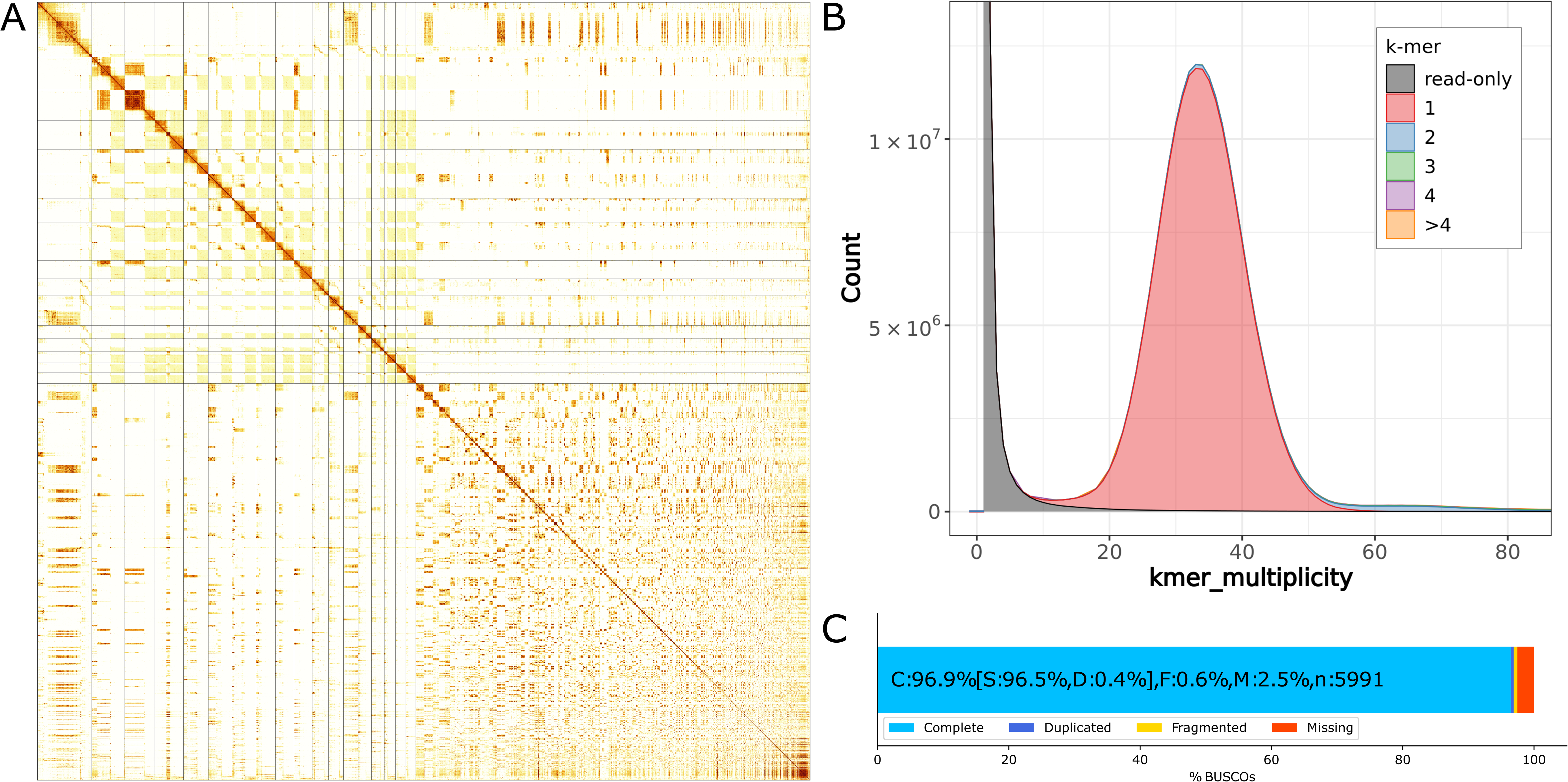
iyXylViol4 assembly of the *Xylocopa violacea* genome. **A)** Hi-C contact map (Supp Methods). Scaffolds are ordered by size with the 17 pseudo-chromosomal super scaffolds appearing in the top left half of the map, defined by overlayed lines. Visualisation constructed with multimapping reads (MAPQ=0). **B)** Merqury kmer spectra, k = 19, single peak representing the haploid male genome of iyXylViol4. **C)** Completeness of the hymenoptera_odb10 BUSCO set (5991 genes).

Following Wallberg *et al*. (2019), we identified the centromeric signature of low GC% in 6 super scaffolds (Supplementary Methods, Figure S5). We identified one such region at the centre of the only firmly identified metacentric chromosome (iyXylViol4_SUPER_4). The other 5 candidates all separate putative euchromatic regions bearing many coding annotations, from regions of high repeat content. This pattern of repeat expansion around centromeric sequences has been observed in other bees, such as *Austroplebeia australis* (Travenzoli *et al*., 2022), and may help to explain the high levels of interaction between unplaced scaffolds and the pseudo-chromosomal units in the iyXylViol4 assembly.

Highly acrocentric karyotypes are well represented within the Xylocopinae, the genus *Ceratina* exhibits species with karyotypes representing 14-17 chromosomes, with ratios of acrocentric to metacentric chromosomes varying between 16:1, 15:2, and 12:5 (Hoshiba and Imai, 1993; Cunha *et al*., 2021). Such patterns are also common in other, more evolutionarily distant bees: *Austroplebeia australis* has been shown to have 14 largely heterochromatic chromosome pairs and four that are fully euchromatic (Travenzoli *et al*., 2022).

Without further investigation, potentially employing ultra-long read technologies, it is not possible to differentiate between N=16 or N=17 from the iyXylViol4 assembly.

### Assembly QC

BUSCO analysis of the iyXylViol4 assembly showed that it contains 96.5% of the 5991 hymenoptra_odb10 set as complete genes, with only 0.4% complete and duplicated, 0.6% fragmented, and 2.5% missing (Figure 2, Table S2). The genic content was not impacted by the scaffolding process as the same metrics are recovered in the contig, scaffolded, and manually curated assemblies. The iyXylViol4 assembly is QV 63.3 and has a kmer completeness of 98.8% (Table S3).

The iyXylViol4 assembly is 1.02Gb in length. Although this is not outside of the upper limits for known genome sizes from the Apidae (e.g. *Melipona capixaba* 1.38Gb, (Tavares *et al*., 2010; Cunha *et al*., 2021), k-mer based estimation of genome size from iyXylViol4 suggests the genome size to be 672 Mb (Table S4, Figure S4). This estimation is in line with the only prediction from the genus *Xylocopa* comes from Ardila-Garcia *et al*. (2010), who report an estimated genome size of 0.69pg (∼675 Mb) for *Xylocopa virginica krombein*. This species is a member of the North American subgenus *Xylocopoides*, thought to have diverged from the genus *Xylocopa*s.*l.* some 34 mya (Leys *et al*., 2002), and so using this estimate as a cross validation for the iyXylViol4 assembly may not be relevant. The 17 pseudo-chromosomal iyXylViol4 super scaffolds (including unloc) are 481.4 Mb in length, representing a large majority of the predicted genome size. As complete reconstruction of the iyXylViol4 chromosomes was not feasible in this study, we have included all unplaced scaffolds in the final assembly, as these likely encompass the remaining genomic content.

### Repeat Content

The majority of the iyXylViol4 assembly was masked as repetitive sequence (821.28 Mb, 80.47%) (Table S5). The predominant category was unclassified repeats, with 755.96 Mb (74.08%). This pattern is consistent with pseudo-acrocentric chromosomes with extremely elongated heterochromatic arms which are frequently observed in bees and wasps (Hoshiba and Imai, 1993). These have been suggested to be induced by saltatory growth of constitutive heterochromatin after centric fission (Hoshiba and Imai, 1993). Bees from the Apinae genus *Melipona* have recently been shown to exhibit up to 73% heterochromatin content (Pereira *et al*., 2021). As is seen in iyXylViol4, bees from the genus *Melipona* also have terminal euchromatic regions (Piccoli *et al*., 2018) which is consistent with the pseudo-acrocentric chromosomal topology derived from *X. appendiculata* (Hoshiba and Imai, 1993), with many chromosomes representing large expansions of heterochromatin repeats around the centromere.

Classification of the repeats within the iyXylViol4 assembly showed the ten most abundant satellite repeat units identified by srf (Zhang *et al*., 2023) to occupy 105.6Mb of the assembly (Table S6). Further decomposition of the satellite repeats present in the iyXylViol4 assembly, using TRASH (Wlodzimierz *et al*., 2023), revealed the predominant monomeric repeat unit to be a 109mer (Figure S7, Figure S8, Table S7). This 109mer or a 217mer (approximately double its length) were highly abundant throughout the putative acrocentric chromosomes (Figure S8) and was repeated with high identity (Figure S7).

We also observe that the putative centromeric sequences are flanked by a distinct repeat signature. In the metacentric iyXylVio4_SUPER_4, the putative centromere has expansions of a 95mer on either side of it. Regions abundant in this 95mer are also seen in 13 of the 16 putative acrocentric pseudo-chromosomal molecules (Figure S8), and these often occur in proximity to the location of the regions of low GC% which are putatively centromeric.

Recent studies have shown telomeric repeat motifs in Hymenoptera to be diverse, including complex telomeric layering resulting from numerous site specific retrotransposon insertions (Lukhtanov, 2022; Zhou *et al*., 2022). The iyXylViol4 assembly shows that *X. violacea* has telomeres enriched for the canonical 5bp ancestral arthropod repeat motif (TTAGG) (Figure S5). The iyXylViol4 assembly also shows that *X. violacea* has varying sub-telomeric repeat sequences, consistent with ‘Type 2’ telomeres suggested by (Lukhtanov and Pazhenkova, 2023) (Figure S6).

### Annotation

The iyXylViol4_EIv1.0 annotation of the iyXylViol4 assembly contains 10,152 high confidence, protein-coding gene models, coding for 26,577 transcripts (Table S8). This number of annotations is well within the range of those generated for contemporary genome assemblies (Table S9). Using the hymenoptera_odb10 database, this annotation represents 99.75% BUSCO completeness at the protein level, with only 34 BUSCO genes duplicated, 3 fragmented and 12 missing (Table S3). The annotation contains an average of 2.49 transcripts per gene, with a mean transcript cDNA size of 3,238.2bp (Table S10). The distribution of coding genes is skewed to the distal end of the 16 pseudo-chromosomal super-scaffolds with putative pseudo-acrocentric structure (Figure S5), supporting the previously suggested topology of highly repetitive pseudo-acrocentric chromosomes expected in *Xylocopa* species (Hoshiba and Imai, 1993; Gokhman, 2023).

## Conclusion

Here, we present a pseudo-chromosomal genome assembly of the Violet Carpenter bee, *Xylocopa violacea*. At 1.02 Gb, the assembly is larger than the predicted genome size (672 Mb), but also represents large regions of highly repetitive, putatively heterochromatic, sequence. Such chromosomal architecture is in line with the small amount of karyotypic resources from the genus and is also supported by the iyXylViol4_EIv1 annotation. <u>The repetitive regions we describe are predominantly made up of 109 and 217mers</u>. The annotated assembly we present fills an important taxonomic gap in the genomic resource set representing Hymenoptera and will also provide a genomic basis for future interpretation of the expanding range of this charismatic and economically important species.

## Acknowledgments

The authors acknowledge Fiona Fraser (Earlham Institute, Norwich) and Michael Quail (Wellcome Sanger Institute, Hinxton) for valuable conversations and advice in developing Hi-C library preparation. The authors would also like to acknowledge the GRiT (Wellcome Sanger Institute, Hinxton), particularly Jo Wood, Tom Mathers, Dominic Absolon, Camilla Santos, Michael Paulini for invaluable mentorship in Hi-C scaffolding and curation. The authors also acknowledge Kamil Hepak (Norwich Bioscience Institutes, Scientific Computing) for significant HPC support.

## Author Contributions

Language used to describe roles below uses the CRediT Taxonomy (credit.niso.org).

AV acted as ERGA sample ambassador, and with NV and BvRM, initiated the **Conceptualisation** of this study; WJN, SMcT, KG, and WH designed the sequencing strategy, assembly of the genome, and all analyses. WJN conducted **Data Curation** throughout the project; GK and DS curated data during genome annotation; DK, AP, and FS curated data for ENA upload through COPO. WJN conducted all **Formal Analysis** outside of genome annotation, which was conducted by GK and DS. **Funding acquisition** was conducted by AV, SMcT, and WH. Primary **Investigation** was conducted by WJN; NI Prepared IsoSeq libraries and Illumina RNA-Seq libraries; TB Sequenced Illumina RNA-seq libraries, PacBio IsoSeq libraries, and PacBio low-input HiFi libraries; AMa prepared Hi-C libraries. AD Developed and improved the Omega EZNA Total RNA extraction protocol **Methodology**; NI Developed and improved the low-input HiFi library preparation protocol methodology; WJN and AMa developed and tested the Hi-C library preparation methodology. WJN and SMcT conducted overall **Project administration**; CW and KB Coordinated the project from sample submission to data delivery; AMcC, GF, and AMo Conceptualised and administrated the ERGA Pilot Project. AV and NV delivered **Resources** by collecting the individuals sequenced. KG led the development of resource data production capability for reference-grade assembly and annotation. WJN wrote code to deploy **Software** as part of the genome assembly project; DS and GK developed and deployed the software used for genome annotation FS, DK, and AP developed and maintain the COPO data brokering software. WH contributed **Supervision** to the whole project, KG provided leadership responsibility for nucleic acid extraction, short-read sequencing, and long-read sequencing; CW Provided supervision and oversight of all project management activities; LC Provided supervision and oversight for Illumina RNA-seq library preparation. WJN conducted **Validation** on all assemblies generated, and generated the **Visualisation**s used in the publication. WJN led the **Writing – Original Draft** with contributions from AV, NV, SMcT and WH. All authors contributed to **Writing – review & editing** of the final manuscript.

