## Supplementary Materials for "The genome sequence of the Violet Carpenter Bee, *Xylocopa violacea* (Linnaeus, 1785): a hymenopteran species undergoing range expansion"

**SUPPLEMENTARY INFORMATION**

**Authors:** Will J Nash[1*], Angela Man[1], Seanna McTaggart[1], Kendall Baker[1], Tom Barker[1], Leah Catchpole[1], Alex Durrant[1], Karim Gharbi[1], Naomi Irish[1], Gemy Kaithakottil[1], Debby Ku[1], Aaliyah Providence[1], Felix Shaw[1], David Swarbreck[1], Chris Watkins[1], Ann M. McCartney[2], Guilio Formenti[3,4], Alice Mouton[4,5,6], Noel Vella[7], Björn M von Reumont[8,9], Adriana Vella[7*], Wilfried Haerty[1,10*]
* Joint Corresponding Authors

**Author Affiliations**
1. The Earlham Institute, Norwich Research Park, Colney Lane, Norwich, NR4 7UZ, UK
2. Genomics Institute, University of California, Santa Cruz, CA 95060, USA
3. The Vertebrate Genome Laboratory, The Rockefeller University, 1240 York Ave, 10065 New York, USA
4. Department of Biology, University of Florence, Sesto Fiorentino, Italy
5. InBios - Conservation Genetics Laboratory, University of Liege, Chemin de la Vallee 4, 4000 Liege, Belgium
6. SEED - Departement des sciences et gestion de l'environnement, University of Liege, Chemin de la Vallee 4, 4000 Liege, Belgium
7. Conservation Biology Research Group, Biology Department, University of Malta, Msida, MSD 2080, Malta
8. LOEWE Center for Translational Biodiversity Genomics (LOEWE-TBG), Senckenberganlage 25, 60325 Frankfurt, Germany
9. Applied Bioinformatics Group, Faculty of Biological Sciences, Goethe University Frankfurt, Max-von-Laue-Str. 13, 60438 Frankfurt, Germany
10. School of Biological Sciences, The University of East Anglia, Norwich, NR4 7TJ, UK

**Authors for Correspondence:**
- Will Nash, The Earlham Institute, Norwich Research Park, Colney Lane, Norwich, NR4 7UZ, +44 (0) 1603 450 974,
- Wilfried Hearty, The Earlham Institute, Norwich Research Park, Colney Lane, Norwich, NR4 7UZ, +44 (0) 1603 450 974,
- Adriana Vella, Conservation Biology Research Group, Biology Department, University of Malta, Msida, MSD 2080, Malta, +356 2340 2790,

Table of Contents

**SUPPLEMENTARY METHODS** 3

SAMPLE COLLECTION 3
HMW DNA EXTRACTION 3
LOW INPUT HIFI LIBRARY PREPARATION 3
RNA EXTRACTION METHODS 3
RNA-SEQ LIBRARY PREPARATION METHODS 4
PACBIO ISO-SEQ LIBRARY PREPARATION AND SEQUENCING 4
ARIMA HIGH COVERAGE HI-C LIBRARY PREPARATION 5
HI-C SEQUENCING METHODS 6
*1. MiSeq 6*
*2. NovaSeq 6*
GENOME ASSEMBLY METHODS 6
MANUAL CURATION METHODS 6
EVIDENCE-BASED GENOME ANNOTATION 7
*1. Repeat identification 7*
*2. Reference guided transcriptome reconstruction 7*
*3. Cross-species protein alignment 7*
*4. Evidence guided gene prediction 7*
*5. Gene calling with Deep Neural Networks 8*
*6. Gene model consolidation 8*
*7. Functional annotation 9*
PLOTTING METHODS 11
*1. Visualisation of Xylocopa violacea Records from GBIF 11*
*2. Visualisation of Hi-C contact map 11*
*3. Visualisation of BUSCO Results 11*
*4. Visualisation of Telomeric Repeats & Super Scaffolds 11*

**SUPPLEMENTARY RESULTS** 12

FEMTO PULSE TRACES 12
ARIMA HIGH COVERAGE HI-C LIBRARY 13
*1. Shearing 13*
*2. Library Preparation 13*
FULL ANNOTATION 14
HI-C MAPPINGS 15
SUPPLEMENTARY FIGURE 4: GENOME SIZE ESTIMATION 15
SUPPLEMENTARY FIGURE 5. VISUALISATION OF SUPER SCAFFOLDS 16
SUPPLEMENTARY FIGURE 6. VISUALISATION OF TELOMERIC REPEAT DIVERSITY 25
SUPPLEMENTARY FIGURE 7. PERCENT IDENTITY, SATELLITE REPEAT, AND MONOMER DISTRIBUTION ON IYXYLVIO4_SUPER_1 26
SUPPLEMENTARY FIGURE 8. REPEAT CONTENT OF PSEUDO-CHROMOSOMAL SCAFFOLDS 27

### Supplementary Methods

#### Sample Collection

The samples used in this study were two individuals drawn from seven collected at the location described in the main text. Four of these were females (iyXylViol2, iyXylViol3, iyXylViol6, iyXylViol8), and the other three were males (iyXylViol4, iyXylViol5, iyXylViol7). iyXylViol2 (ERS10526492) was used to generate RNA-seq & PacBio Iso-Seq. iyXylViol4 (ERS10526494) was used to generate PacBio HiFi and Hi-C data. iyXylViol5 (ERS10526495) was used in the development of the protocol for the Hi-C data. All other individuals were returned to the ERGA Sample Ambassador (Prof A. Vella, University of Malta, corresponding author) upon completion of the project.

#### HMW DNA Extraction

DNA concentration was measured using the Qubit® 1X dsDNA HS kit (Invitrogen®, Q33230) and the Nanodrop® One Microvolume UV-Vis Spectrophotometer (Thermo Scientific®, ND-ONE-W) was used to measure extraction purity. The distribution of HMW DNA fragment sizes was measured using the FEMTO Pulse® System (Agilent, P/N M5330AA).

#### Low Input HiFi Library Preparation

A low input library was prepared from a starting input of 406 ng gDNA. The sample was sheared using the Megaruptor 3 instrument (Diagenode, P/N B06010003), 4 ng/µl at a speed 33. The sample underwent AMPure® PB bead (PacBio®, P/N 100-265-900) purification and concentration at a ratio of 0.45X before undergoing library construction using SMRTbell template prep kit 2.0 (PacBio®, P/N 100-983-900).

The HiFi library was prepared from 284 ng of sheared gDNA. Using SMRTbell® Express Template Prep Kit 2.0 (PacBio®, P/N 100-983-900), a barcoded adapter (PacBio®, P/N 101-628-500) was ligated to create the SMRTbell library. The final SMRTbell library was AMPure purified initially with 0.45X ratio and again with AMPure beads at 35% v/v with Elution Buffer (PacBio®, P/N 101-633-500) 3.1X diluted beads to library. Final library size was estimated from a smear analysis performed on the FEMTO Pulse® System (Agilent, P/N M5330AA) and quantified by fluorescence (Invitrogen Qubit™ 3.0, P/N Q33216).

The loading calculations for sequencing were completed using the PacBio® SMRT®Link Binding Calculator v10.2.0.133424. Sequencing primer v4 was annealed to the adapter sequence of the library. The library binding to the sequencing polymerase was completed using Sequel® II Binding Kit v2.0 (PacBio®, P/N 101-842-900). Calculations for primer to template and polymerase to template binding ratios were kept at default values for the library type. Sequel® II DNA internal control was spiked into the library complex at the standard concentration prior to sequencing. Sequencing was performed using Sequel® II Sequencing Plate 2.0 (PacBio®, P/N 101-820-200) chemistry and Instrument Control Software v10.1.0.125432.

#### RNA Extraction Methods

For each extraction, 15 mg of the tissue of interest was collected from the frozen specimen and placed in a 2 ml Eppendorf tube to which a 5 mm tungsten bead and 700 uL of lysis buffer had already been added. The manufacturer recommendation when working with <20 mg material is to use 350uL lysis buffer, but 700 uL was used here to facilitate optimal separation of the debris from the supernatant following the first downstream centrifuge step. For the head and thorax samples, 20 uL/ml of 2-mercaptoethanol was added to the lysis buffer prior to homogenization; this modification was a response to issues encountered during amplification using previously extracted RNA from these tissues.

The samples were then processed for 5 minutes using a SPEX SamplePrep GenoGrinder set to 1000 rpm, and the homogenate was spun at 16,000 g for 5 minutes on a tabletop centrifuge. The supernatant was transferred to a 1.5 ml Eppendorf tube, and 1x volume of 100% ethanol was added. The manufacturer protocol recommends using 70% ethanol, but 100% ethanol was used here to maximise preservation of small RNAs. From here the remainder of the extraction procedure followed the handbook guide (V4.0, August 2019) for animal tissue extraction, including the on-column DNase treatment.

As a final post-extraction precaution, the RNA samples for the head and thorax extractions were cleaned up using a sodium acetate precipitation. This lowered the RNA yield by approximately 60% but improved the purity readings (as measured by the Nanodrop).

#### RNA-seq Library Preparation methods

1 ug of total RNA was purified to extract mRNA with a Poly(A) mRNA Magnetic Isolation Module. The technology is based on the coupling of Oligo d(T)25 to paramagnetic beads which facilitates the binding of poly(A)+ RNA. Isolated mRNA was then fragmented for 12 minutes at 94°C, and first strand cDNA was synthesised. This process reverse transcribes the RNA fragments primed with random hexamers into first strand cDNA using reverse transcriptase and random primers. The second strand synthesis process removes the RNA template and synthesizes a replacement strand to generate ds cDNA. Directionality is retained by adding dUTP during the second strand synthesis step and subsequent cleavage of the uridine containing strand using USER Enzyme (a combination of UDG and Endo VIII). NEBNext Adaptors were ligated to end-repaired, dA-tailed DNA. The NEBNext Adaptors with novel hairpin loop structure are designed to ligate with high efficiency and minimize adaptor-dimer formation. The loop contains a U, which is removed by treatment with USER Enzyme to open the loop and make it available as a substrate for PCR. The ligated products were subjected to a bead-based purification using Beckman Coulter AMPure XP beads (A63882) to remove most of un-ligated adaptors. Adaptor Ligated DNA was then enriched by receiving 10 cycles of PCR (30 secs at 98°C, 10 cycles of: 10 secs at 98°C _75 secs at 65°C _5 mins at 65°C, final hold at 4°C). Barcodes are incorporated during PCR using NEBNext Multiplex Oligos for Illumina® (96 Unique Dual Index Primer Pairs) thereby allowing multiplexing. The quality of the resulting libraries was determined using Agilent High Sensitivity DNA Kit from Agilent Technologies (5067-4626) and the concentration measured with a High Sensitivity Qubit assay from ThermoFisher (Q32854).

The library pool was diluted down to 0.5 nM using EB (10 mM Tris pH 8.0) in a volume of 18 ul before spiking in 1% Illumina phiX Control v3 (FC-110-3001). This was denatured by adding 4 ul 0.2N NaOH and incubating at room temperature for 8 mins, after which it was neutralised by adding 5 ul 400 mM tris pH 8.0. A master mix of EPX1, EPX2, and EPX3 from Illumina's NovaSeq Xp 2-lane kit v1.5 (20043130) was made and 63 ul added to the denatured pool leaving 90 ul at a concentration of 100 pM. This was loaded onto a NovaSeq SP flow cell using the NovaSeq Xp Flow Cell Dock. The flow cell was then loaded onto the NovaSeq 6000 along with an NovaSeq 6000 SP cluster cartridge, buffer cartridge, and 300 cycle SBS cartridge (20028400). The NovaSeq had NVCS v1.7.5 and RTA v3.4.4 and was set up to sequence 150 bp PE reads. The data was demultiplexed and converted to fastq using bcl2fastq2.

#### PacBio Iso-Seq Library Preparation and Sequencing

PacBio Iso-Seq libraries were constructed starting from 234-300 ng of total RNA per sample. Reverse transcription cDNA synthesis was performed using NEBNext® Single Cell/Low Input cDNA Synthesis & Amplification Module (NEB, E6421). Each cDNA sample was amplified with barcoded primers for a total of 12 cycles. The barcoded cDNA samples were pooled equimolar before SMRTbell library construction. The library pool was prepared according to the guidelines laid out in the IsoSeq protocol version 02 (PacBio, 101-763-800), using SMRTbell express template prep kit 2.0 (PacBio, 102-088-900). The library pool was quantified using a Qubit Fluorometer 3.0 (Invitrogen) and sized using the Bioanalyzer HS DNA chip (Agilent Technologies, Inc.).

The loading calculations for the Iso-Seq library pool using the PacBio SMRTlink Binding Calculator v10.2.0.133424 and prepared for sequencing applicable to the library type. Sequencing primer v4 was annealed to the Iso-Seq library pool and complexed to the sequencing polymerase with the Sequel II binding kit V2.1 (PacBio, 101-843-000). Calculations for primer to template and polymerase to template binding ratios were kept at default values for the library type. Sequencing internal control complex 1.0 (PacBio, 101-717-600) was spiked into the final complex preparation at a standard concentration before sequencing for all preparations. The sequencing chemistry used was Sequel® II Sequencing Plate 2.0 (PacBio®, 101-820-200) and the Instrument Control Software v10.1.0.125432.

The Iso-Seq pool was sequenced on the Sequel II instrument with one Sequel II SMRT®cell 8M cell. The parameters for sequencing were diffusion loading, 30-hour movie, 2-hour immobilisation time, 2-hour pre-extension time, 60 pM on plate loading concentration.

#### Arima High Coverage Hi-C Library Preparation

Hi-C library preparation was conducted following the Arima High Coverage Hi-C Kit (A410110), following the kit protocol (A160162 v01). The process was conducted using a pellet of 25% of the head tissue form the same male *Xylocopa violacea* individual (iyXylViol4) from which thorax tissue was used to generate PacBio HiFi reads for this assembly (see main text).

AMPure XP Beads (A63881) were used as the DNA Purification Beads throughout.

Cross linking was conducted using the Low Input protocol (p15-16). Following estimation of the input amount, the Arima High Coverage HiC Protocol (p19-22) was conducted with the following modifications:

- Step 17 - Proximally ligated DNA was eluted in **55 µl** of Elution Buffer instead of 100 µl. This is due to the requirements of the Covaris instrument used for DNA fragmentation.

The Arima QC1 step was passed, and the sample was taken through the initial steps of the Arima-HiC 2.0 Library Preparation using Swift Biosciences Accel-NGS 2S Plus DNA Library Kit (A160164 v00). DNA fragmentation was conducted using the Covaris ML230 focused-ultrasonicator, with an ML230_500661 Rack 8 microTUBE strip (50 + 1.8 mm offset). The settings used on the Covaris instrument are descried in Table S12. Following DNA shearing and QC, DNA size selection was conducted using AMPure XP Beads. Following the Arima protocol, this step was followed by Biotin enrichment with the following modification:

- Step 14 - modified by the addition of **50 µl** on Elution Buffer instead of 40 µl. This is due to the volumes required for the NEB Next Ultra II DNA Library Prep Kit for Illumina.

Following Biotin Enrichment, the NEBNext Ultra II DNA Library Prep Kit for Illumina (E7645S) was used to prepare the sequencing library from the End Repair step onwards. The kit manual (version 7.0_9/22) was followed with the modifications described here:

- An additional AMPure XP bead clean-up (0.8x) was conducted to remove excess adaptor prior to PCR-enrichment of adapter-ligated DNA (NEB Protocol Step 4.1). This clean-up did not follow the standard AMPure XP bead clean-up protocol as, at this point, the library is mounted on streptavidin beads. For this clean-up step, the magnetic pellet is **not discarded**, and the following elution step (AMPure XP Protocol (B37419AB.book, p4), Step 5). The 'mixed bead' solution was then cleaned up by a repeat of steps 5-14 of the previously employed Biotin enrichment protocol (Arima-HiC 2.0 Library Preparation using Swift Biosciences Accel-NGS 2S Plus DNA Library Kit Protocol (A160164 v00), Section 2.3, p8-9). This was done to remove the maximum amount of AMPure XP beads whilst retaining the highest concentration of library. Advice received following the completion of this project suggests that this clean-up step was not necessary, and it would be possible to conduct the PCR Amplification with AMPure XP beads present. This was not done in this study.
- At step 4.1.1, NEBNext Ultra II Q5 master mix was replaced with an equal volume of KAPA Hifi Hotstart Readymix (KK2601).
- At step 4.1.3, PCR was conducted for 8 cycles, where the NEB protocol recommendation for the input amount (NEB Protocol Table 4.1) suggested 6 cycles.
- Library was constructed using NEBNext® Multiplex Oligos for Illumina (E7335S) Index 11 (E7321AVIAL).

#### Hi-C Sequencing Methods

Prior to the full sequencing run used in the assembly, a quality control sequencing run was conducted using the Illumina MiSeq Instrument. This was done to check that the library contained sufficient contact information to be used for Hi-C Scaffolding.

##### 1. MiSeq

The pools were diluted to 2 nM and denatured using 2N NaOH before diluting to 20 pM with Illumina HT1 buffer. The denatured pool was loaded on an Illumina MiSeq Sequencer with a 300 cycle MiSeq Reagent Micro Kit v2 (Illumina MS-103-1002) at 10 pM concentration with a 1% phiX control v3 spike (Illumina FC-110-3001) as per Illumina's recommendations with MiSeq Control Software version 4.0.0.1769 and RTA version 1.18.54.4. The data was demultiplexed and converted to fastq using bcl2fastq2.

##### 2. NovaSeq

The library pool was diluted down to 0.5 nM using EB (10 mM Tris pH 8.0) in a volume of 18 ul before spiking in 1% Illumina phiX Control v3 (FC-110-3001). This was denatured by adding 4 ul 0.2N NaOH and incubating at room temperature for 8 mins, after which it was neutralised by adding 5 ul 400mM tris pH 8.0. A master mix of EPX1, EPX2, and EPX3 from Illumina's NovaSeq Xp 2-lane kit v1.5 (20043130) was made and 63 ul added to the denatured pool leaving 90ul at a concentration of 100 pM. This was loaded onto a NovaSeq SP flow cell using the NovaSeq Xp Flow Cell Dock. The flow cell was then loaded onto the NovaSeq 6000 along with an NovaSeq 6000 SP cluster cartridge, buffer cartridge, and 300 cycle SBS cartridge (20028400). The NovaSeq had NVCS v1.7.5 and RTA v3.4.4 and was set up to sequence 150 bp PE reads. The data was demultiplexed and converted to fastq using bcl2fastq2.

#### Genome Assembly Methods

As the sequenced individual was haploid, hifiasm was run without duplication purging (`-l 0`). Following assessment of the initial kmer plot produced by hifiasm, the coverage threshold for homozygous reads was manually tuned (`--hom-cov 33`).

Hi-C Reads were trimmed with trimmomatic (v0.39, (Bolger, Lohse, and Usadel 2014) settings: `ILLUMINACLIP:adapters.fa:2:30:10 SLIDINGWINDOW:4:15 MINLEN:70`

Assembly size statistics were calculated using abyss-fac (v2.3.5, (Jackman et al. 2017)).

Following manual curation, the initial PacBio HiFi long reads used for assembly were mapped back to the curated assembly to assess final coverage, this was done using minimap2 (v2.24, (Li 2018)) with the settings `-ax map-hifi -t 128`, piping the output to samtools (v1.15.1, (Li et al. 2009)) to generate a bam file, using the settings `-@ 127 -o ${OUTPUT}.bam --write-index -`

#### Manual Curation Methods

Following GRiT curation, the curated assembly was uploaded to the ERGA Nextcloud along with the raw data involved in its generation and was re-evaluated by the GRiT Team. Following this step, the final manually curated assembly was downloaded from the ERGA Nextcloud and passed forward to Annotation. Scaffolds joined during manual curation were labelled SUPER, to denote super scaffolds, or pseudochromosoal units. Where scaffolds could be associated with a pseudochromosoal unit but finalised positions could not be resolved, a numbered '_unloc_' suffix was added to the header. Final curation of the sequence files and headers was conducted using seqkit (v2.5.1, (Shen et al. 2016)).

#### Evidence-based genome annotation

##### 1. Repeat identification

Repeat annotation was performed using the EI-Repeat version 1.3.4 pipeline (https://github.com/EI-CoreBioinformatics/eirepeat) which uses third party tools for repeat calling. In the pipeline, RepeatModeler (v1.0.11 - http://www.repeatmasker.org/RepeatModeler/) was used for de novo identification of repetitive elements from the assembled *Xylocopa* genome. Unclassified repeats were searched in a custom BLAST database of organellar genomes (mitochondrial sequences from *Apis mellifera*, OK075087.1). Any repeat families matching organellar DNA were also hard-masked. Repeat identification was completed by running RepeatMasker v4.0.72 with a RepBase Aculeata library and with the customized RepeatModeler library (i.e. after masking out protein coding genes), both using the -nolow setting.

##### 2. Reference guided transcriptome reconstruction

Gene models were derived from the RNA-Seq, IsoSeq transcripts (HQ+LQ) and Full-Length Non-Concatamer Reads (FLNC) using the REAT transcriptome workflow (https://github.com/EI-CoreBioinformatics/reat). HISAT2 v 2.2.1 (Kim et al., 2019) was selected as the short read aligner with IsoSeq transcripts aligned with minimap2 v 2.18-r1015 (Li H., 2018), maximum intron length was set as 100,000 bp and minimum intron length to 20bp. IsoSeq alignments were required to meet 95% coverage and 90% identity. High-confidence splice junctions were identified by Portcullis v 1.2.4 Mapleson, et al. 2018). RNA-Seq Illumina reads were assembled for each tissue, as well as for a pooled 'all' dataset that contained reads from all tissues and the additional reads downloaded from SRA (SRR14690757, (Koludarov et al. 2023), with StringTie2 v2.1.5 (Kovaka et al., 2019) (Table S13) and Scallop v v0.10.5 (Shao and Kingsford, 2017) (Table S14), while FLNC reads were assembled using StringTie2 v2.1.5 (Table S15). Gene models were derived from the RNA-Seq assemblies and IsoSeq and FLNC alignments with Mikado (https://github.com/EI-CoreBioinformatics/mikado, Venturini et al., 2018). Mikado was run with all Scallop, StringTie2, IsoSeq and FLNC alignments and a second run with only IsoSeq and FLNC alignments.

##### 3. Cross-species protein alignment

Protein sequences from 17 Apidae species (Table S16) were aligned to the Carpenter Bee assembly using the REAT Homology workflow (https://github.com/EI-CoreBioinformatics/reat) with options –annotation_filters aa_len –alignment_species InsectAg –filter_max_intron 60000 –filter_min_exon 10 –alignment_filters aa_len internal_stop intron_len exon_len splicing –alignment_min_coverage 90 –junction_f1_filter 40 –post_alignment_clip clip_term_intron-exon –term5i_len 15000 –term3i_len 15000 –term5c_len 36 –term3c_len 36. The REAT Homology workflow aligns proteins with spaln v2.4.7 (Gotoh et al., 2008) and filters and generates metrics to remove misaligned proteins. Simultaneously, the same protein set were also aligned using miniprot v0.3 (Li, 2023) and similarly filtered as in the REAT homology workflow. The aligned proteins from both methods were clustered into loci and a consolidated set of gene models were derived via Mikado.

##### 4. Evidence guided gene prediction

Evidence guided annotation of protein coding genes was carried out using the REAT prediction workflow making use of the repeat annotation, RNA-Seq alignments, transcript assembly and alignment of protein sequences. The pipeline has four main steps:

1. The REAT transcriptome and homology Mikado models are categorised based on alignments to Uniprot Apoidea proteins (downloaded 01-09-23) to identify models with likely full-length CDS and which meet basic structural checks i.e., having complete but not excessively long UTR's and not exceeding a minimum CDS/cDNA ratio. A subset of gene models is then selected from the classified models and used to train the AUGUSTUS gene predictor (Stanke et al., 2005).
2. Augustus is run with extrinsic evidence generated in the REAT transcriptome and homology runs (repeats, protein alignments, RNA-Seq alignments, splice junctions, categorised Mikado models). Three evidence guided AUGUSTUS predictions is created using alternative bonus scores and priority based on evidence type.

3. AUGUSTUS models, REAT transcriptome / homology models, protein and transcriptome alignments are provided to EVidenceModeler (EVM) (Haas et al., 2008) to generate consensus gene structures.
4. EVM models are processed through Mikado to add UTR features and splice variants.

Results are summarised in Table S17.

#### 5. Gene calling with Deep Neural Networks

Genes were predicted using Helixer a deep neural networks (DNN) approach with the publicly available invertebrate model with options `--subsequence-length 213840 --overlap-offset 106920 --overlap-core-length 160380`

Results are summarised in Table S18.

#### 6. Gene model consolidation

The final set of gene models was selected using Minos (https://github.com/EI-CoreBioinformatics/minos). Minos is a pipeline that generates and utilises metrics derived from protein, transcript, and expression data sets to create a consolidated set of gene models. In this annotation, the following gene were filtered and consolidated into a single set of gene models using minos:

1) The three alternative evidence guided Augustus gene builds derived from the REAT prediction run described earlier.
2) The gene models derived from the REAT transcriptome runs described earlier.
3) The gene models derived from the REAT homology run described earlier.
4) The gene models derived from Helixer.

Gene models were classified as biotypes protein_coding_gene, predicted_gene and transposable_element_gene, and assigned as high or low confidence based on the criteria below:

a) **High confidence protein_coding_gene**: Any protein coding gene where any of its associated gene models have a BUSCO v5.4.7 (Seppey et al., 2019) protein status of Complete/Duplicated OR have diamond (v0.9.36) coverage (average across query and target coverage) >= 90% against the listed protein datasets mentioned in Section 3 or Uniprot Apoidea proteins. Or alternatively have average blastp coverage (across query and target coverage) >= 80% against the list protein datasets / Uniprot Apoidea AND have transcript alignment F1 score (average across nucleotide, exon and junction F1 scores based on RNA-Seq transcript assemblies) >= 60%.

b) **Low confidence protein_coding_gene**: Any protein coding gene where all of its associated transcript models do not meet the criteria to be considered as high confidence protein coding transcripts.

c) **High confidence transposable_element_gene**: Any protein coding gene where any of its associated gene models have coverage >= 40% against the combined interspersed repeats (see repeat identification methods section 1). These annotations are not included in the iyXylViol_EIv1.0 annotation.

d) **Low confidence transposable_element_gene**: Any protein coding gene where all of its associated transcript models do not meet the criteria to be considered as high confidence and assigned as a transposable_element_gene (see c). These annotations are not included in the iyXylViol_EIv1.0 annotation.

e) **Low confidence predicted_gene**: Any protein coding gene where all of the associated transcript models do not meet the criteria to be considered as high confidence protein coding transcripts. In addition, where any of the associated gene models have average blastp coverage (across query and target coverage) < 30% against the protein datasets mentioned AND having a protein-coding potential score < 0.25 calculated using CPC2 0.1 (Kong et al., 2007). These annotations are not included in the iyXylViol_EIv1.0 annotation.

f) **Low ncrna gene**: Any gene model with no CDS features AND a protein-coding potential score < 0.25 calculated using CPC2 0.1. These annotations are not included in the iyXylViol_EIv1.0 annotation.

g) **Discarded models**: Any models having no BUSCO protein hit AND a protein alignment score (average across nucleotide, exon and junction F1 scores based on protein alignments) <0.2 AND a transcript alignment F1 score (average across nucleotide, exon and junction F1 scores based on RNA-Seq transcript assemblies) <0.2 AND a diamond coverage (target coverage) <0.3 AND Kallisto v0.44 (Bray et al., 2016) expression score <0.2 from across RNA-Seq reads OR having short CDS <30bps. Any ncRNA genes (no CDS features) not meeting the ncrna gene requirements (f) were also excluded. These annotations are not included in the iyXylViol_EIv1.0 annotation.

#### 7. Functional annotation

All the proteins were annotated using AHRD v.3.3.3 (Hallab et al., 2014; https://github.com/groupschoof/AHRD/blob/master/README.textile). Sequences were blasted against the UniProt Apoidea sequences (data download 01-09-23), both Swiss-Prot and TrEMBL datasets (The UniProt Consortium, 2014). Proteins were BLASTed (v2.6.0; blastp) with an e-value of 1e-5. We also provided interproscan (v5.22.61; Jones et al., 2014) results to AHRD. We adapted the standard AHRD example configuration file path test/resources/ahrd_example_input_go_prediction.yml, distributed with the AHRD tool, changing the following apart from the location of input and output files:

1. We included the GOA mapping from uniprot (ftp://ftp.ebi.ac.uk/pub/databases/GO/goa/UNIPROT/goa_uniprot_all.gaf.gz) as parameter 'gene_ontology_result',
2. We also included the interpro database (ftp://ftp.ebi.ac.uk/pub/databases/interpro/61.0/interpro.xml.gz) and provided as parameter 'interpro_database',
3. We changed the parameter 'prefer_reference_with_go_annos' to 'false'
4. The blast database specific weights used were:

```
blast_dbs:
  swissprot:
    weight: 653
    description_score_bit_score_weight: 2.717061
  trembl:
    weight: 904
    description_score_bit_score_weight: 2.590211
```

### Plotting methods

##### 1. Visualisation of Xylocopa violacea Records from GBIF

The distribution map presented in Figure 1 was plotted from occurrence data downloaded from GBIF (see main text) using python-dwca-reader (https://python-dwca-reader.readthedocs.io/en/latest/) , geopandas-v0.8.1 (Jordahl et al. 2020, http://doi.org/10.5281/zenodo.3946761, https://geopandas.org/ ) and h3-pandas (https://h3-pandas.readthedocs.io/en/latest/) in a custom python script available here:
https://github.com/bibionid/xylocopa_assembly/blob/main/map_from_gbif.py

##### 2. Visualisation of Hi-C contact map

The Hi-C contact map presented in Figure 2 was generated using PretextSnapshot (https://github.com/wtsi-hpag/PretextSnapshot) with the settings: `--sequences "=full" -r 600 -c 23 --gridSize 0`. Lines to demark the super scaffolds were manually overlayed using InkScape v1.2.2 (https://inkscape.org/).

##### 3. Visualisation of BUSCO Results

The BUSCO Results presented in Figure 2 were plotted using a custom python script available here: https://github.com/bibionid/xylocopa_assembly/blob/main/busco_plotter.py

##### 4. Visualisation of Telomeric Repeats & Super Scaffolds

The visualisations of the iyXylViol4 assembly super scaffold structure (Figure S4) and telomeric repeat distribution (Figure S5) were plotted using pyGenomeTracks (https://github.com/deeptools/pyGenomeTracks).

Where Hi-C contact matrices is visualised in these figures, Hi-C data was remapped to the final manually curated iyXylViol4 assembly using `bowtie2` (Langmead and Salzberg 2012). From these mappings, the contact matrix, in .cool format, was generated using hicBuildMatrix from the HiCExplorer Suite (Ramírez et al. 2018, https://github.com/deeptools/HiCExplorer/).

Where HiFi read coverage is visualised in these figures, adapter trimmed HiFi reads were mapped to the final manually curated iyXylViol4 assembly as described in the main text. Output BAM files were used raw (MAPQ=0) and filtered for multimapping reads (MAPQ=10), and were transformed into bedgraph format using bedtools (Quinlan and Hall 2010) genomecov -bg.

Where the proportion of GC content is visualised in these figures, 10kb windows spanning the final manually curated iyXylViol4 assembly were created using `bedtools makewindows`. %GC was calculated in these windows using `bedtools nuc`. The output was transformed into bedgraph format using `grep -v "^#" iyXylViol4_10kbwindows.nuc | cut -f1,2,3,5 > iyXylViol4_10kbwindows_GC.bg`

In Figure S4, telomeric repeats are the output of `tidk search` with the canonical arthropod telomeric repeat (TTAGG) as input. In Figure S5 telomeric repeats are visualised from the output of `tidk explore` run on iyXylViol4_SUPER_1 and iyXylViol4_SUPER_4 respectively. These are plotted from bedgraph format.

Where the iyXylViol4 annotation is visualised in these figures, this is plotted from the annotation in GTF format, with transcripts condensed to a consensus gene model for clarity of visualisation.

### Supplementary Results

#### Femto PULSE Traces

**Supplementary Figure 1. Femto PULSE traces for the two high molecular weight (HMW) DNA samples extracted from male *Xylocopa violacea* (iyXylViol4) thorax tissue.** Two ~30mg sections of thorax tissue were extracted as described in the methods. Two elutions of HMW DNA were taken from the magnetic beads used, both are presented here. All four of these samples were pooled prior to sequencing. **A)** Tissue section 1 Elution 1, **B)** Tissue section 1 Elution 2, **C)** Tissue section 2 Elution 1, **D)** Tissue section 2 Elution 2.

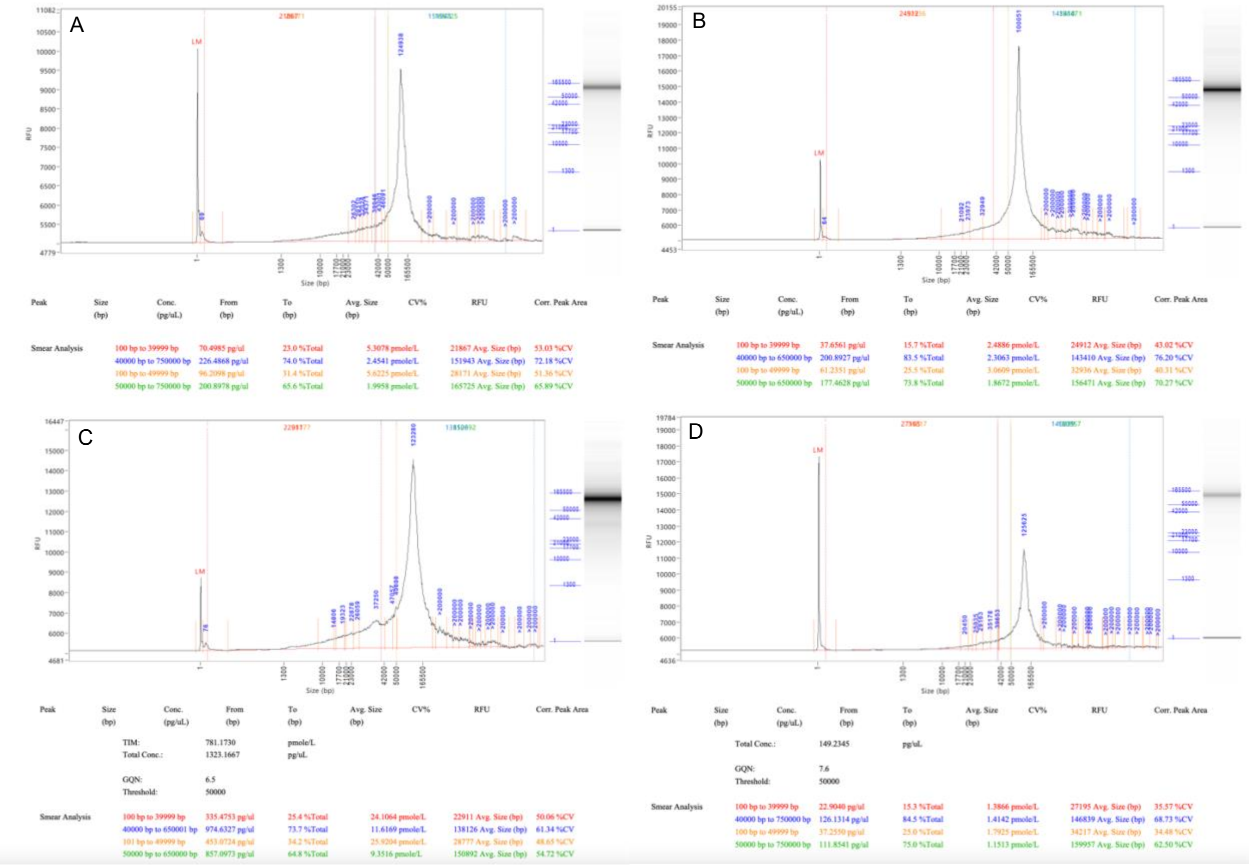

#### Arima High Coverage Hi-C Library

Head Tissue from individual iyXylViol4, was used to generate 14.5 ng/µl of proximity ligated DNA.

In the Arima QC1 step, a Qubit concentration of 9.26 ng/µl suggested a yield of 64.82ng of DNA, and scored 86.4% on the Arima-QC1 Metric, achieving a PASS status.

##### 1. Shearing

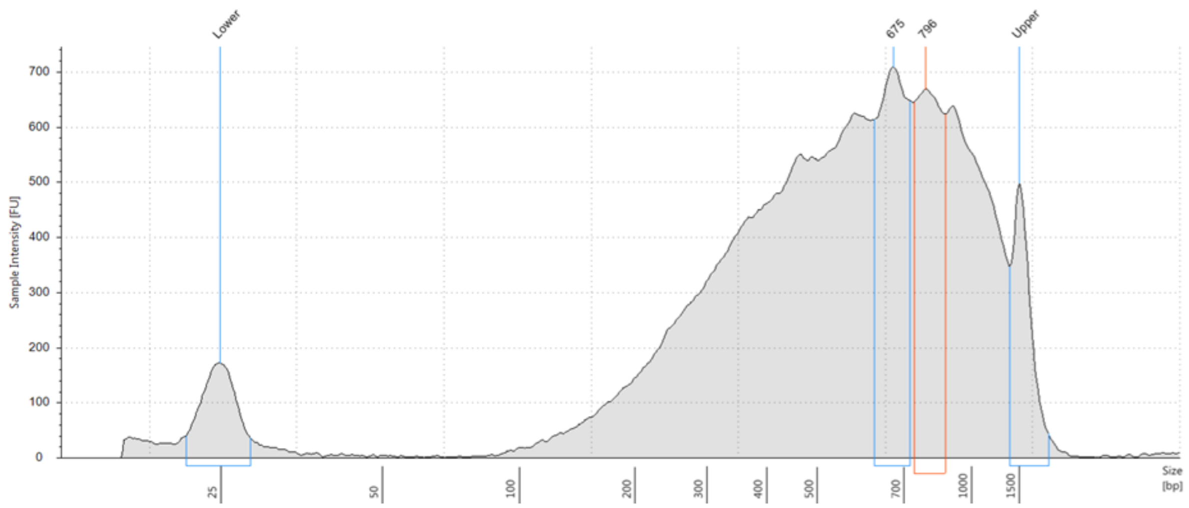


**Supplementary Figure 2.** Fragment distribution of proximity ligated DNA from the head tissue of male *Xylocopa vioalcea* (iyXylViol4), following fragmentation on the COVARIS ML230 focused-ultrasonicator, following the settings and treatment described in Table S12.

Following size selection, 3.4 ng/µl of fragmented, size selected, proximity ligated DNA was generated.

Following biotin enrichment, 2.9 ng/µl of biotin enriched, fragmented, size selected, proximity ligated DNA, at this point mounted on beads was generated.

##### 2. Library Preparation

QC following Adapter ligation showed DNA concentration to be 0.9 ng/µl.

Following the NEBNext Ultra II DNA Library Prep, a library was created with an average library insert size of 720 bp and a predicted molarity of 1.7 nM. Following 8 PCR cycles, this library showed a Qubit concentration of 0.8 ng/µl and showed a fragment distribution as shown in Fig. S3. This library was submitted for sequencing by the Earlham Institute Transformative Genomics team as described in the main text and Supplementary Methods.

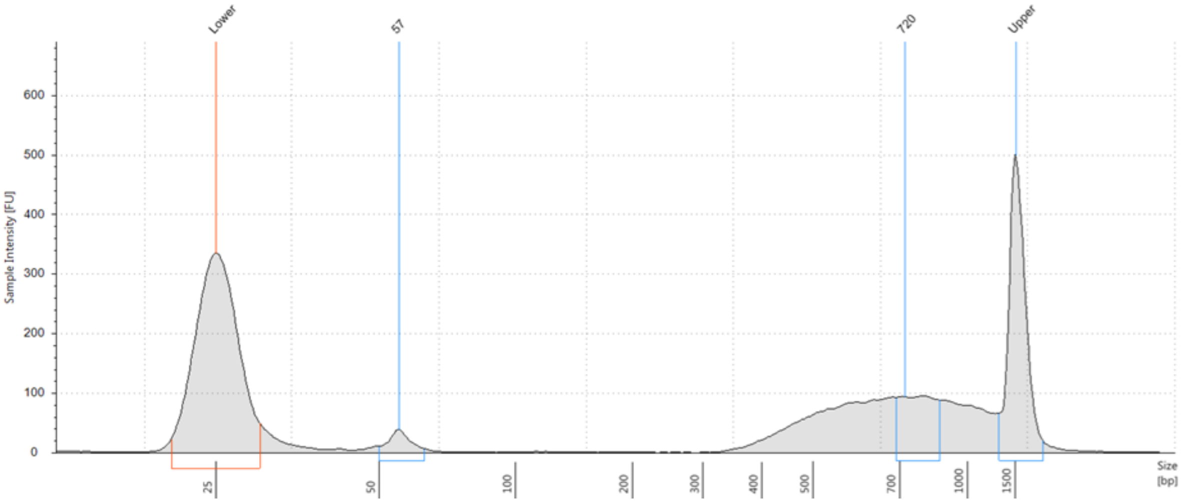


**Supplementary Figure 3.** Fragment distribution of library created from proximity ligated DNA from the head tissue of *Xylocopa vioalcea* (iyXylViol4), using the NEBNext Ultra II DNA Library Prep Kit, following 8 PCR cycles.

#### Full Annotation

The full iyXylViol4_EIv1.0.release.gff3 version of the annotation we generated, containing transposable element gene and ncRNA gene predictions as well as lower confidence predictions and functional predictions is available at: https://zenodo.org/doi/10.5281/zenodo.13221010.

#### Hi-C Mappings

Hi-C reads were mapped to the contig assembly generating 266,533,180 unique mappings (52.3%), leading to 89,151,712 Hi-C contacts, with 75,047,295 and 41,171,942 being intrachromosomal and long range (>20kb) contacts respectively.

#### Supplementary Figure 4: Genome Size Estimation

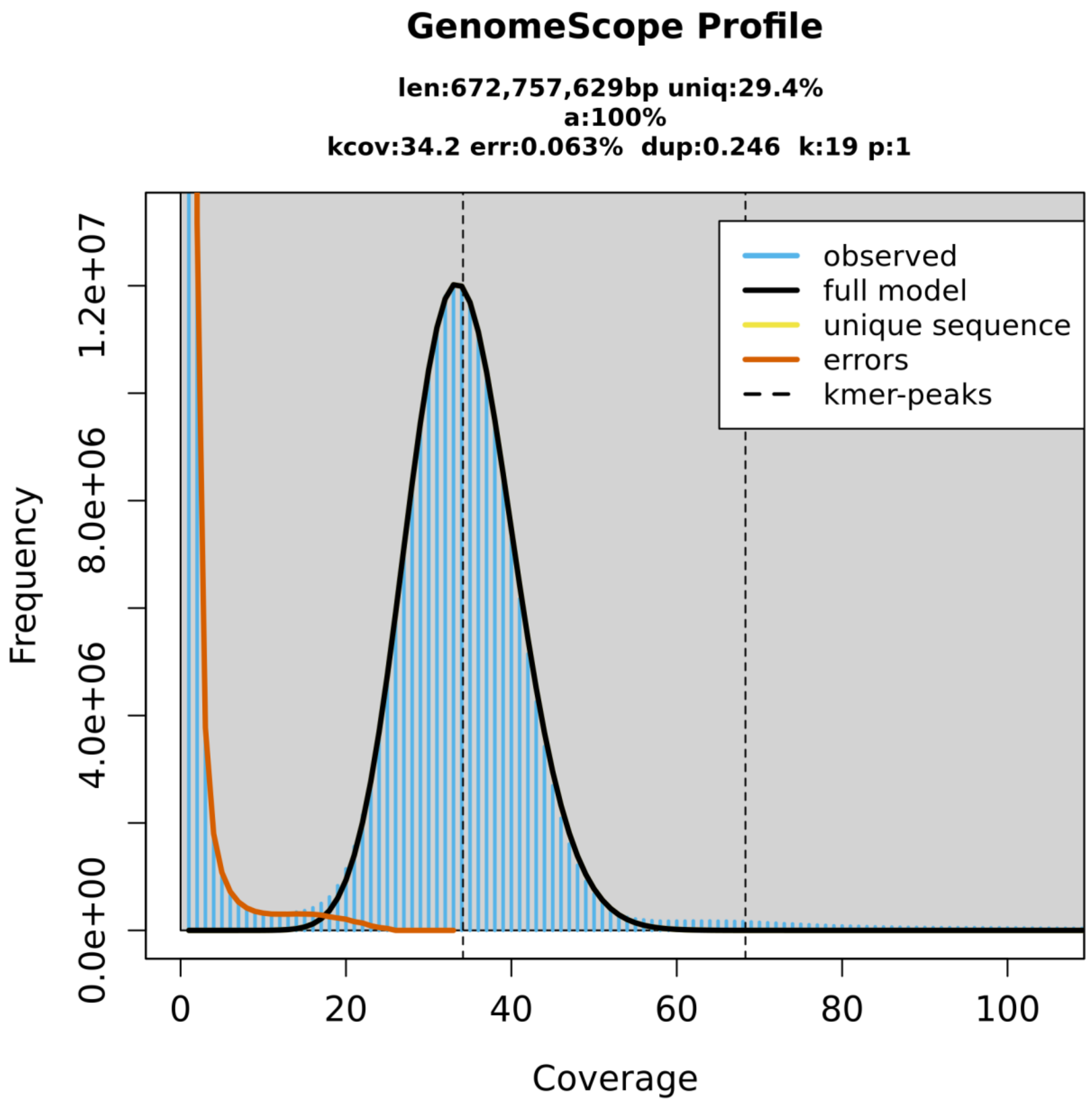


**Supplementary Figure 4. k-mer based genome size estimation of the iyXylViol4 assembly.** Estimation and figure generated using GeneScopeFK (Table S1) with settings p=1 and k=19. Model results are presented in Table S4.

#### Supplementary Figure 5. Visualisation of Super Scaffolds

**Supplementary Figure 5. Super Scaffolds of the iyXylViol4 genome assembly.** The 17 panels below visualise the super scaffolds of the iyXylViol4 assembly of the *Xylocopa violacea* genome. All panels have the same structure: **Top Row:** Hi-C Contact Matrix, Hi-C Reads mapped back to the finalised manually curated iyXylViol4 assembly, **Second Row:** PacBio HiFi Read mapping, HiFi reads used for assembly remapped to the finalised manually curated iyXylViol4 assembly (Green = MAPQ 0, Red = MAPQ 10), **Third Row:** Proportion of the nucleotides within 10Kb windows that are GC. **Fourth Row:** Canonical Arthropod telomeric repeat (TTAGG) density. **Fifth Row:** iyXylViol4_EIv1.0 annotation, transcripts condensed to consensus gene model for clarity. **Putative centromeres**, identified by characteristic low levels of %GC (e.g. (Wallberg et al. 2019)), are highlighted by blue volumes. These are present on super scaffolds 1, 2, 4, 5, 8, & 11.

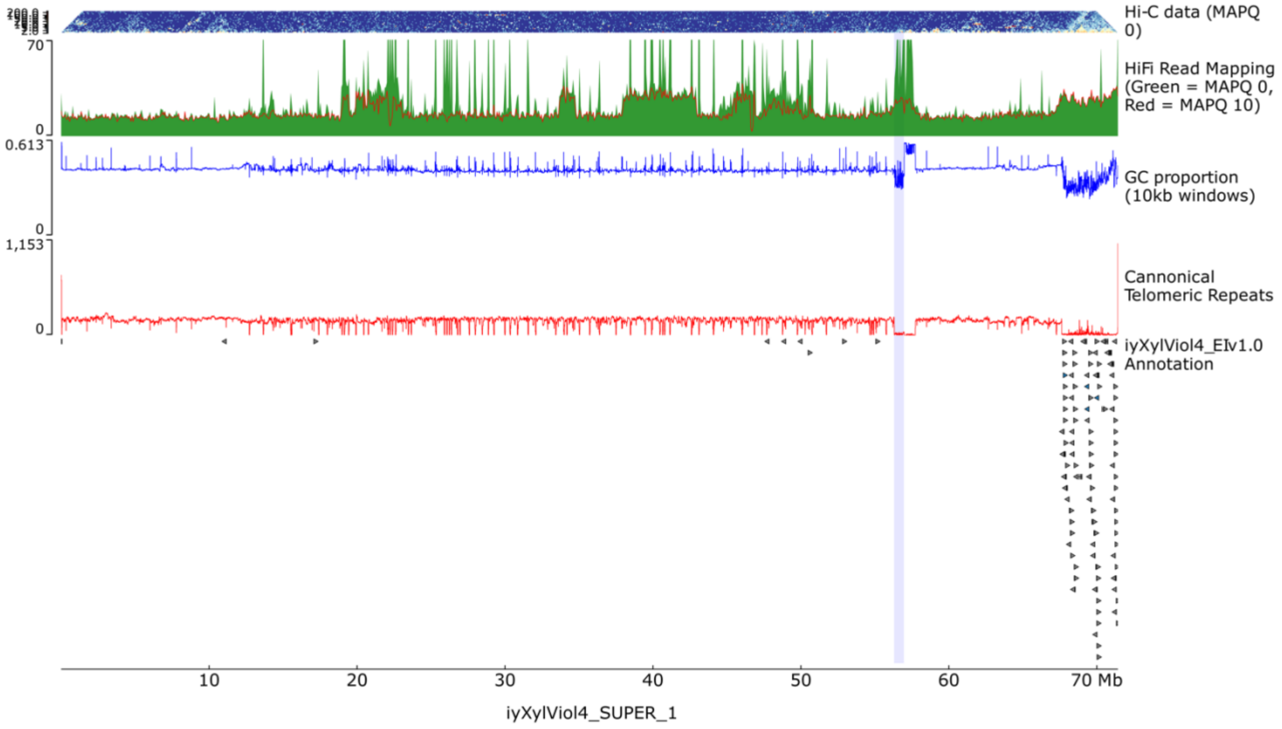


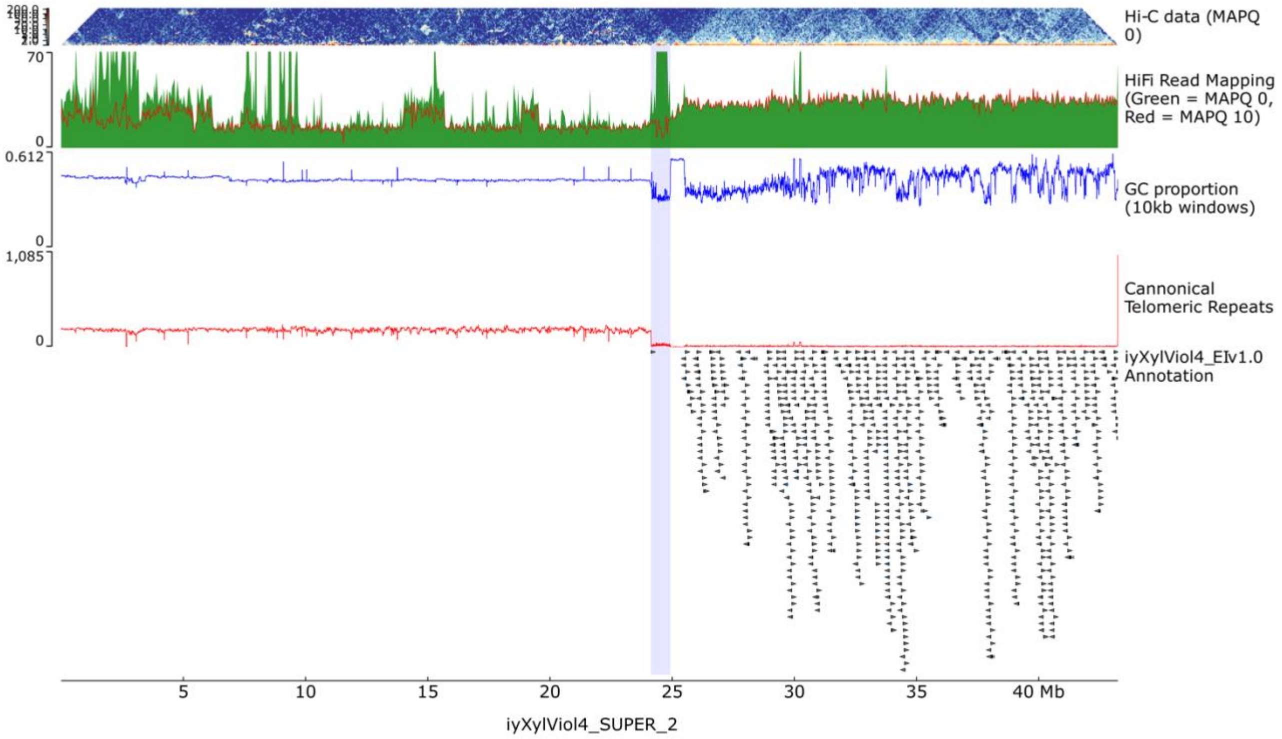

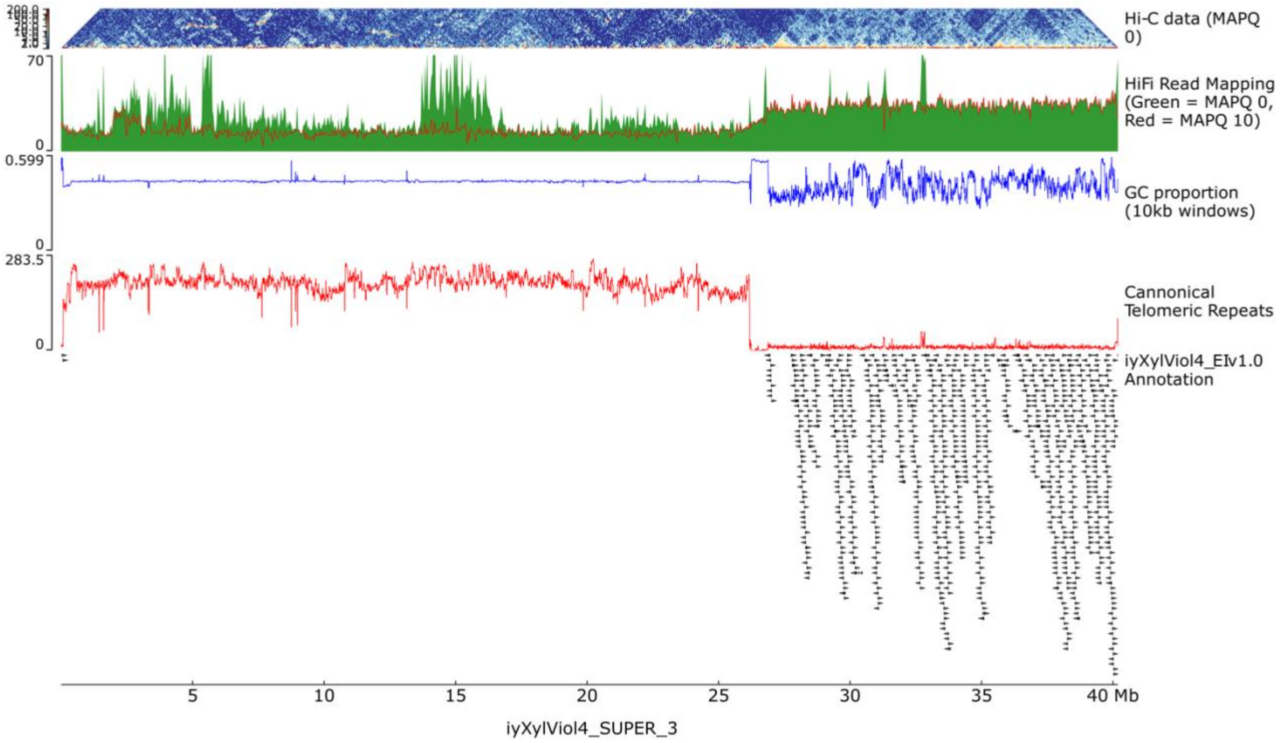
Hi-C data (MAPQ 0)
HiFi Read Mapping (Green = MAPQ 0, Red = MAPQ 10)
GC proportion (10kb windows)
Cannonical Telomeric Repeats
iyXylViol4_EIv1.0 Annotation
70
0
0.599
283.5
5
10
15
20
25
30
35
40 Mb
iyXylViol4_SUPER_3

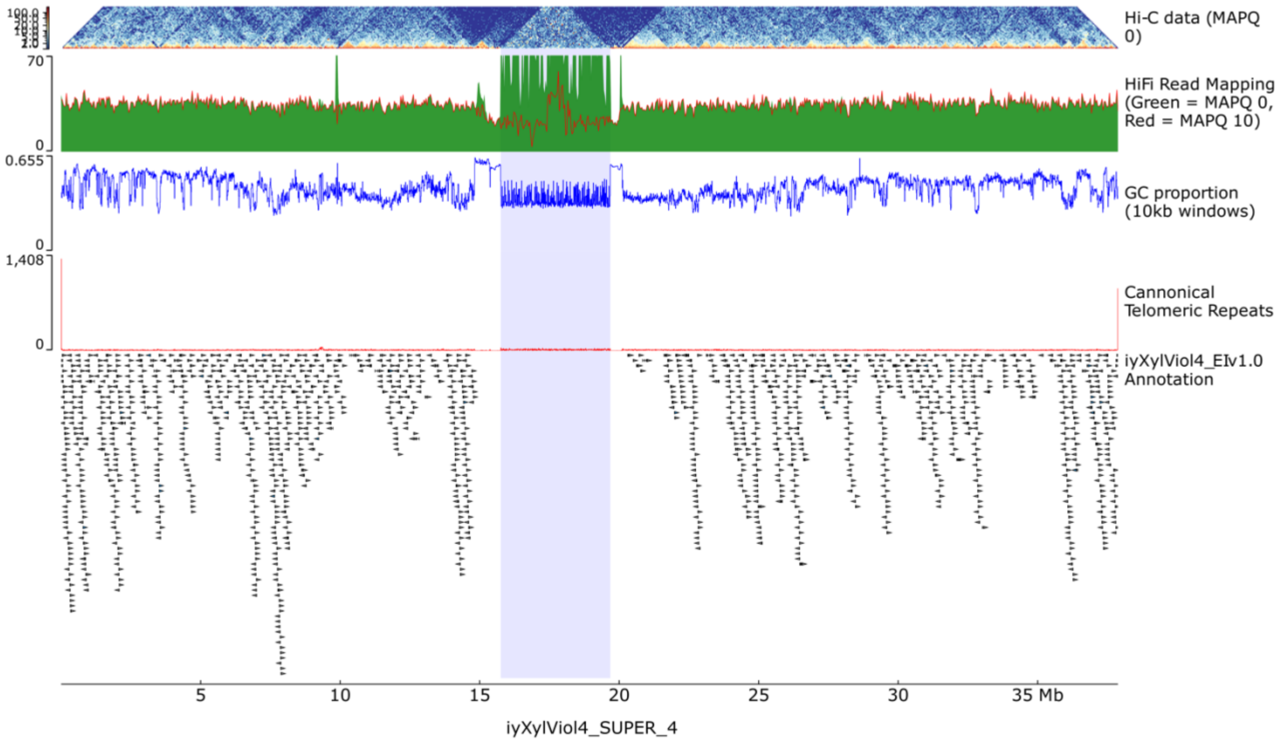
Hi-C data (MAPQ 0)
HiFi Read Mapping (Green = MAPQ 0, Red = MAPQ 10)
GC proportion (10kb windows)
Cannonical Telomeric Repeats
iyXylViol4_EIv1.0 Annotation
70
0
0.655
1,408
5
10
15
20
25
30
35 Mb
iyXylViol4_SUPER_4

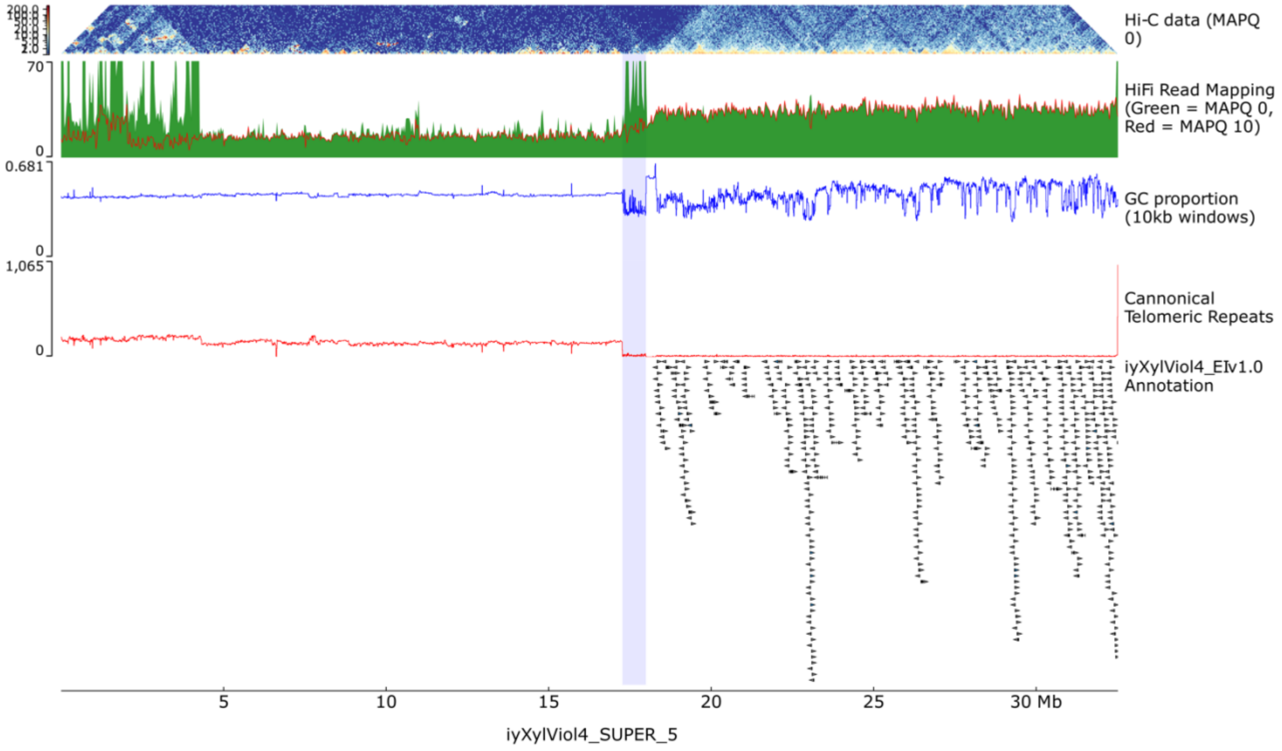
Hi-C data (MAPQ 0)
70
0
HiFi Read Mapping (Green = MAPQ 0, Red = MAPQ 10)
0.681
0
GC proportion (10kb windows)
1,065
0
Cannonical Telomeric Repeats
iyXylViol4_EIv1.0 Annotation
5
10
15
20
25
30 Mb
iyXylViol4_SUPER_5

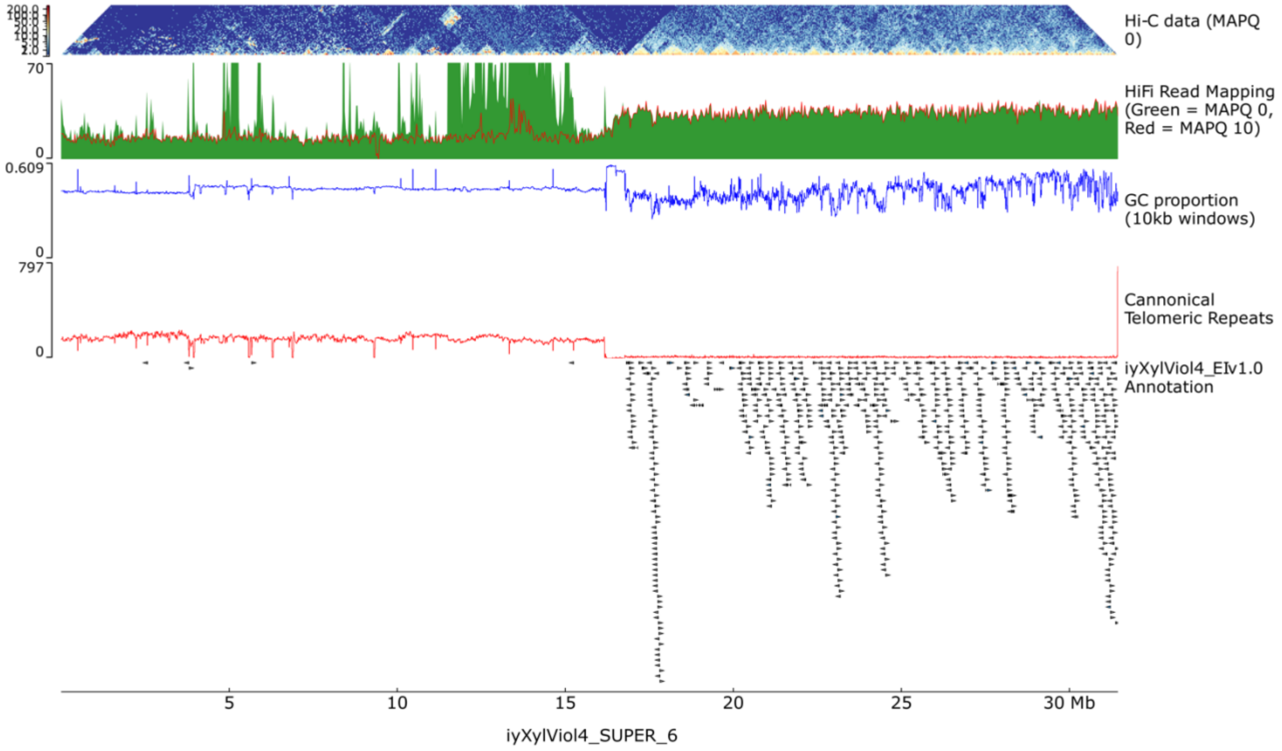
Hi-C data (MAPQ 0)
70
0
HiFi Read Mapping (Green = MAPQ 0, Red = MAPQ 10)
0.609
0
GC proportion (10kb windows)
797
0
Cannonical Telomeric Repeats
iyXylViol4_EIv1.0 Annotation
5
10
15
20
25
30 Mb
iyXylViol4_SUPER_6

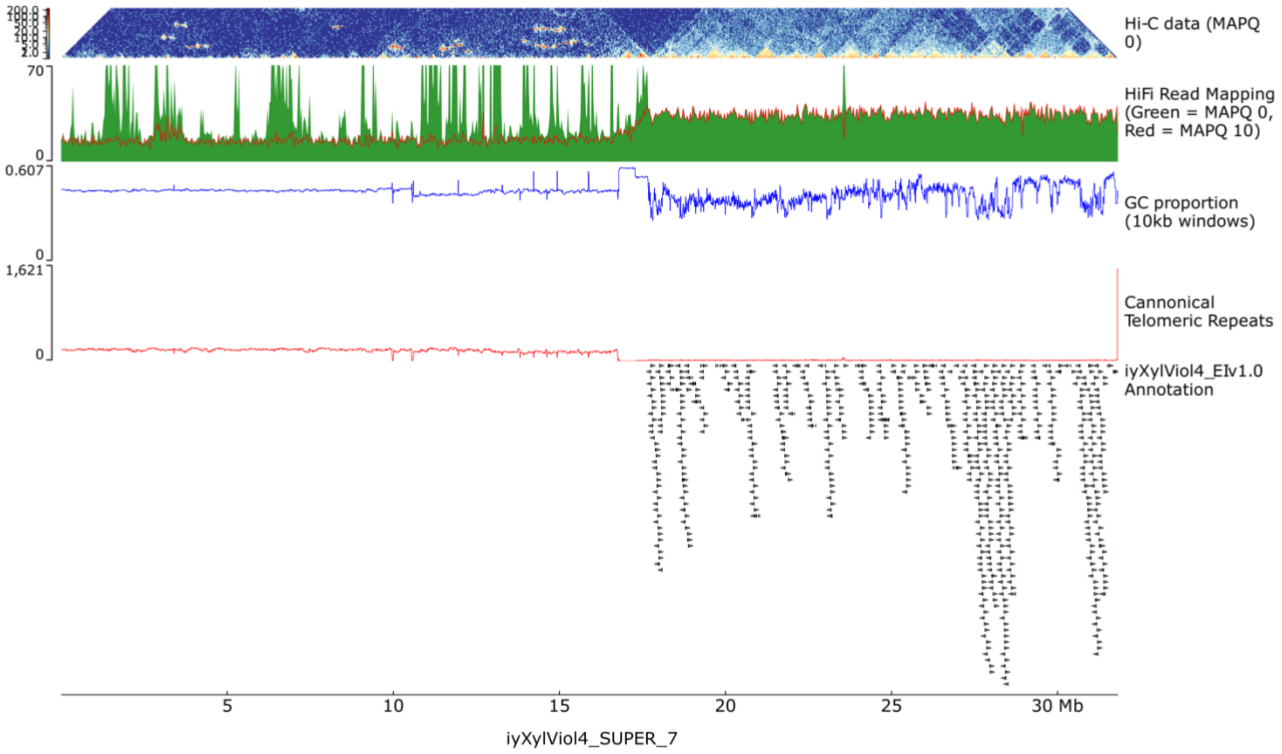

Hi-C data (MAPQ 0)
HiFi Read Mapping (Green = MAPQ 0, Red = MAPQ 10)
GC proportion (10kb windows)
Cannonical Telomeric Repeats
iyXylViol4_EIv1.0 Annotation
70
0
0.607
0
1,621
0
5
10
15
20
25
30 Mb
iyXylViol4_SUPER_7


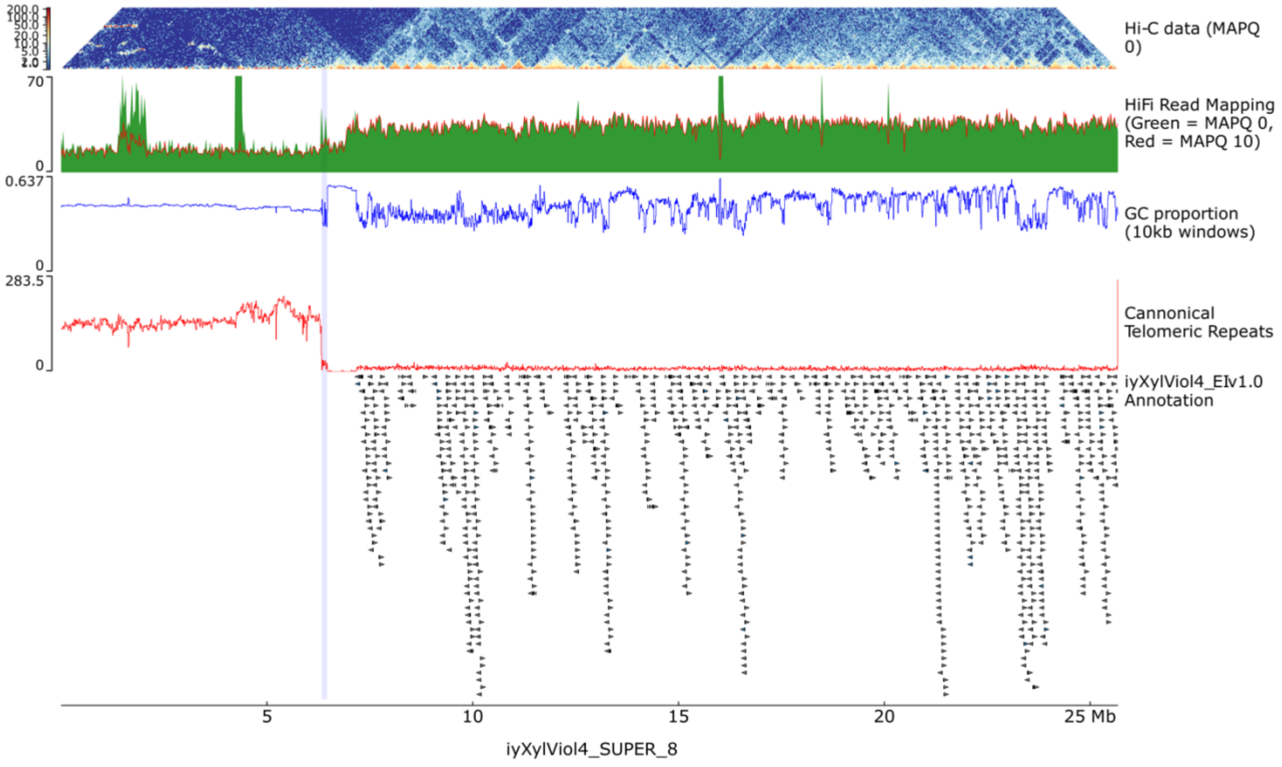

Hi-C data (MAPQ 0)
HiFi Read Mapping (Green = MAPQ 0, Red = MAPQ 10)
GC proportion (10kb windows)
Cannonical Telomeric Repeats
iyXylViol4_EIv1.0 Annotation
70
0
0.637
0
283.5
0
5
10
15
20
25 Mb
iyXylViol4_SUPER_8

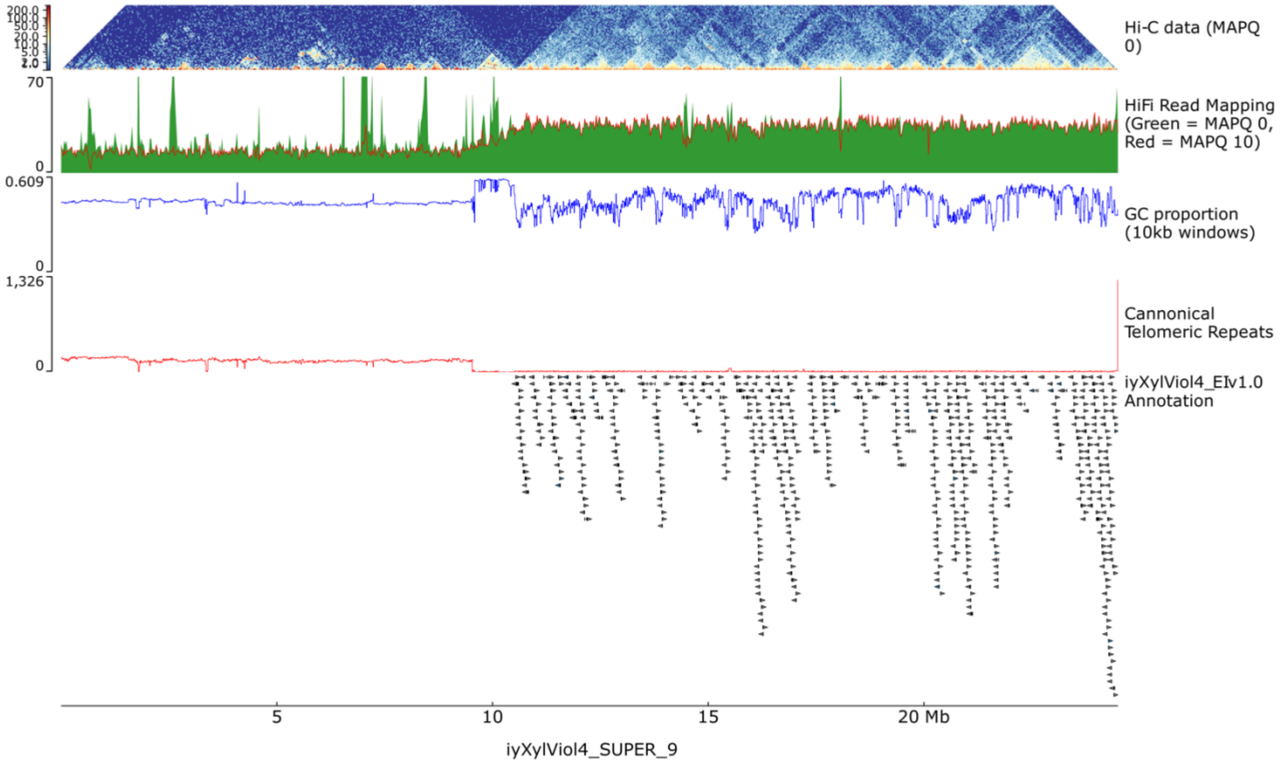

Hi-C data (MAPQ 0)
HiFi Read Mapping (Green = MAPQ 0, Red = MAPQ 10)
GC proportion (10kb windows)
Cannonical Telomeric Repeats
iyXylViol4_EIv1.0 Annotation
70
0
0.609
0
1,326
0
5
10
15
20 Mb
iyXylViol4_SUPER_9


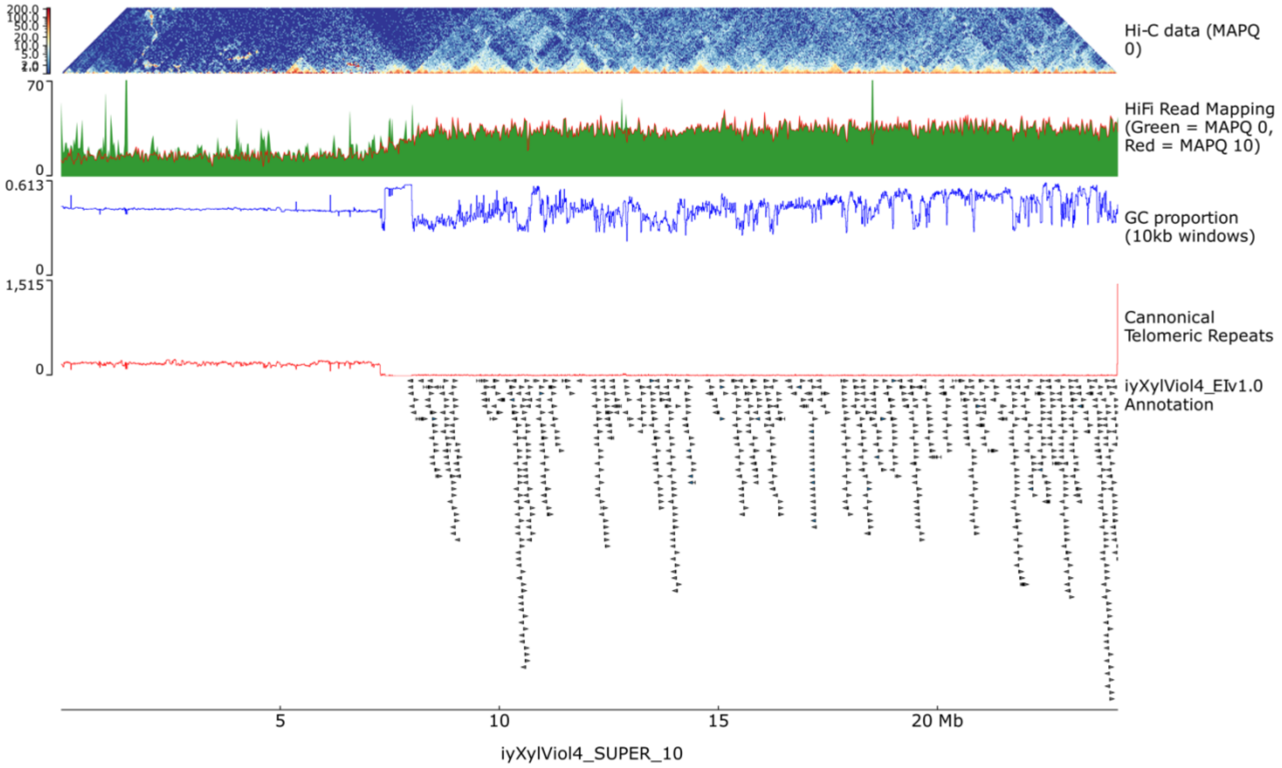

Hi-C data (MAPQ 0)
HiFi Read Mapping (Green = MAPQ 0, Red = MAPQ 10)
GC proportion (10kb windows)
Cannonical Telomeric Repeats
iyXylViol4_EIv1.0 Annotation
70
0
0.613
0
1,515
0
5
10
15
20 Mb
iyXylViol4_SUPER_10

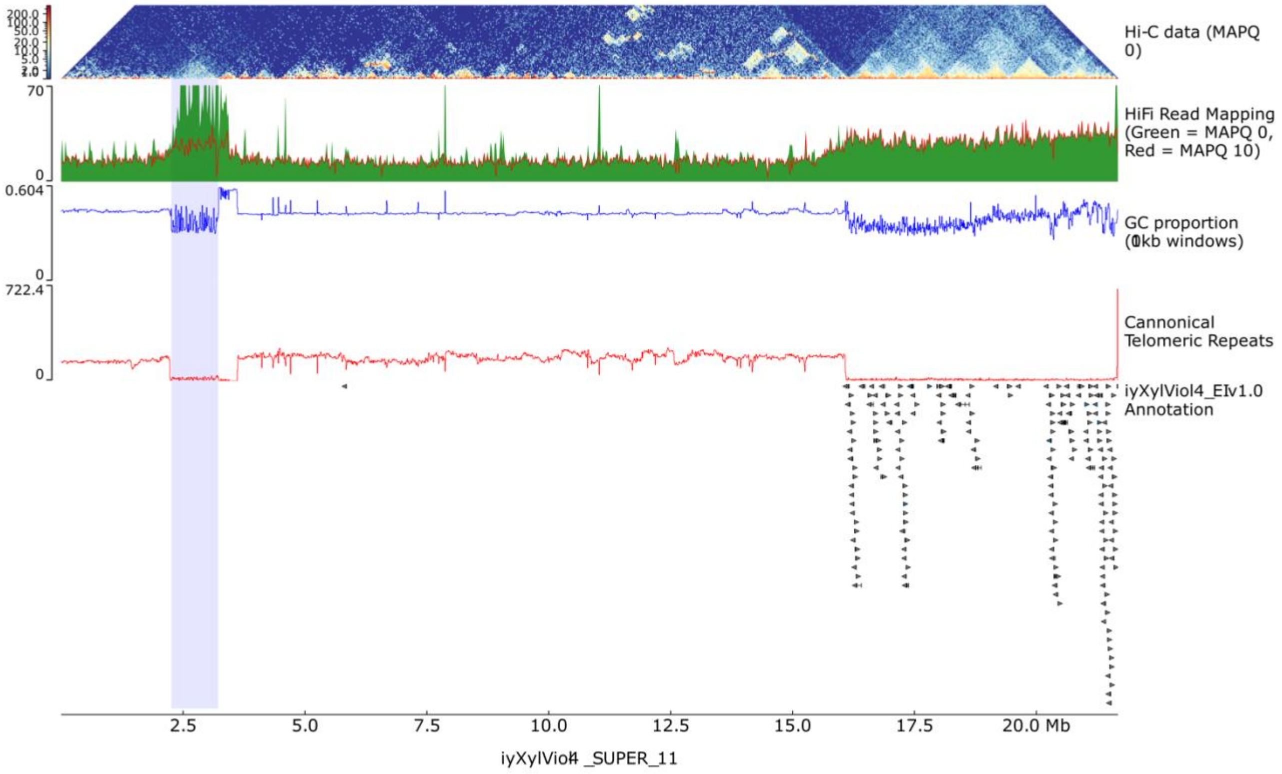
Hi-C data (MAPQ 0)
HiFi Read Mapping (Green = MAPQ 0, Red = MAPQ 10)
GC proportion (10kb windows)
Cannonical Telomeric Repeats
iyXylViol4_EIv1.0 Annotation
200.0
100.0
50.0
20.0
10.0
5.0
2.0
70
0
0.604
0
722.4
0
2.5
5.0
7.5
10.0
12.5
15.0
17.5
20.0 Mb
iyXylViol4 _SUPER_11

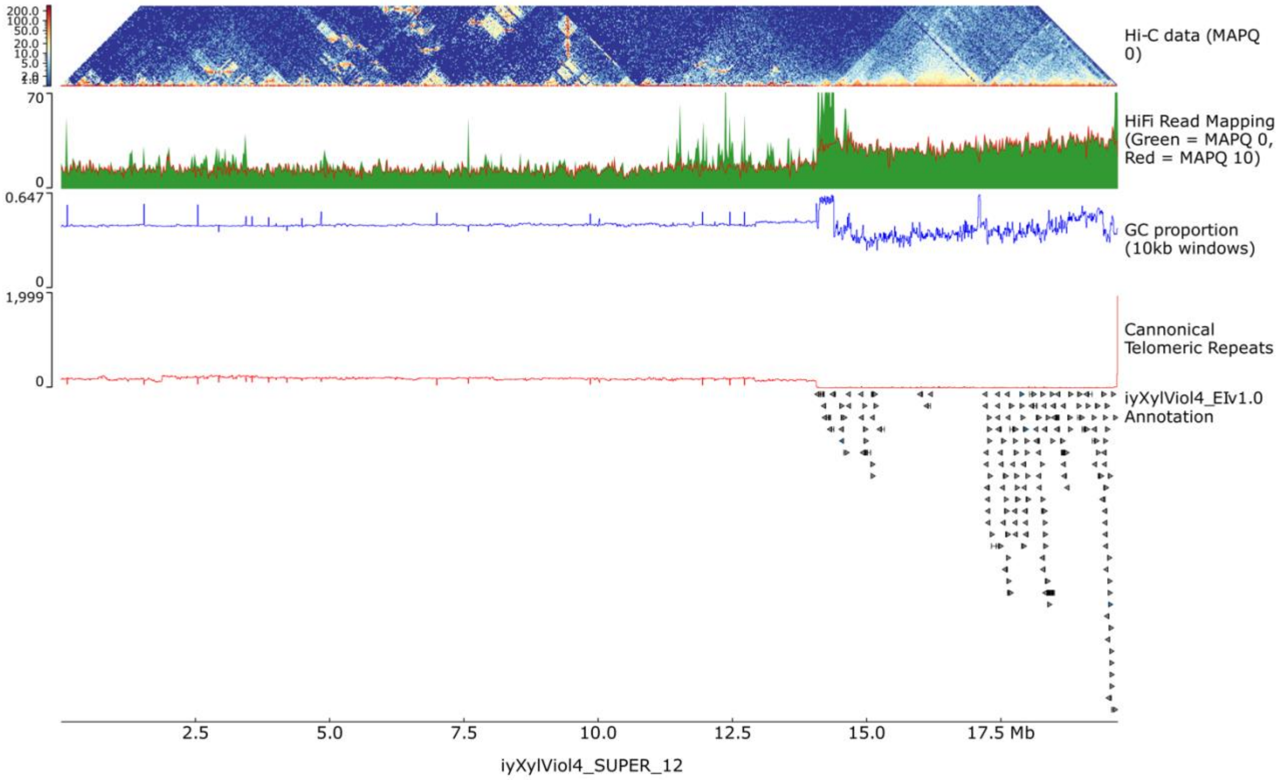
Hi-C data (MAPQ 0)
HiFi Read Mapping (Green = MAPQ 0, Red = MAPQ 10)
GC proportion (10kb windows)
Cannonical Telomeric Repeats
iyXylViol4_EIv1.0 Annotation
200.0
100.0
50.0
20.0
10.0
5.0
2.0
70
0
0.647
0
1,999
0
2.5
5.0
7.5
10.0
12.5
15.0
17.5 Mb
iyXylViol4_SUPER_12

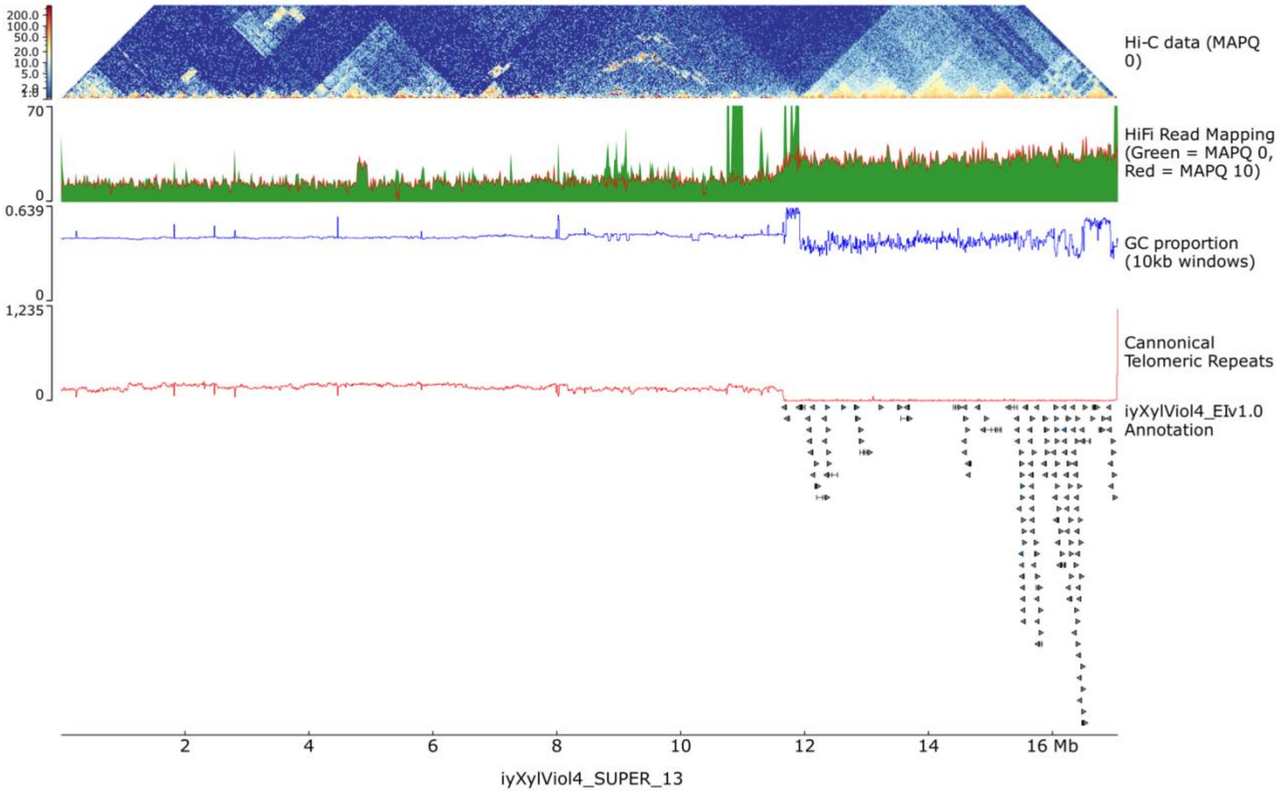
Hi-C data (MAPQ 0)
HiFi Read Mapping (Green = MAPQ 0, Red = MAPQ 10)
GC proportion (10kb windows)
Cannonical Telomeric Repeats
iyXylViol4_EIv1.0 Annotation
iyXylViol4_SUPER_13

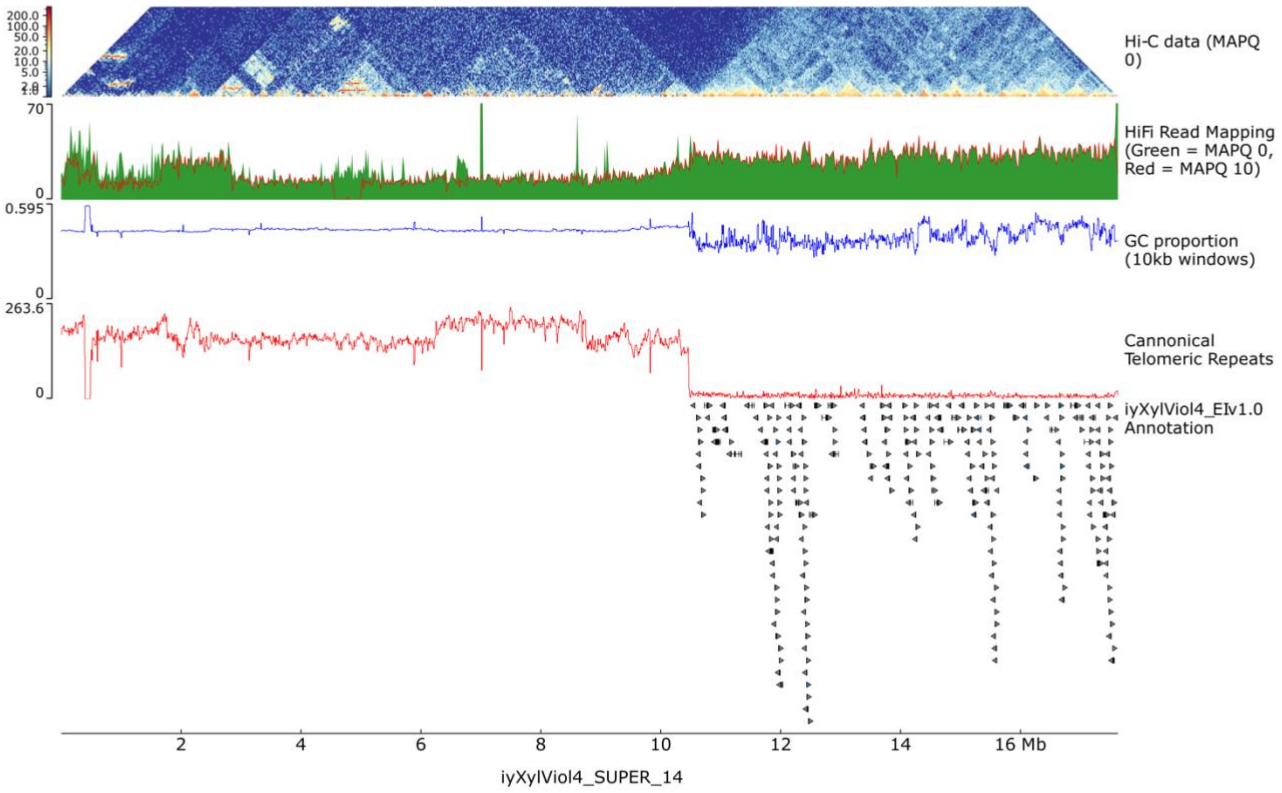
Hi-C data (MAPQ 0)
HiFi Read Mapping (Green = MAPQ 0, Red = MAPQ 10)
GC proportion (10kb windows)
Cannonical Telomeric Repeats
iyXylViol4_EIv1.0 Annotation
iyXylViol4_SUPER_14

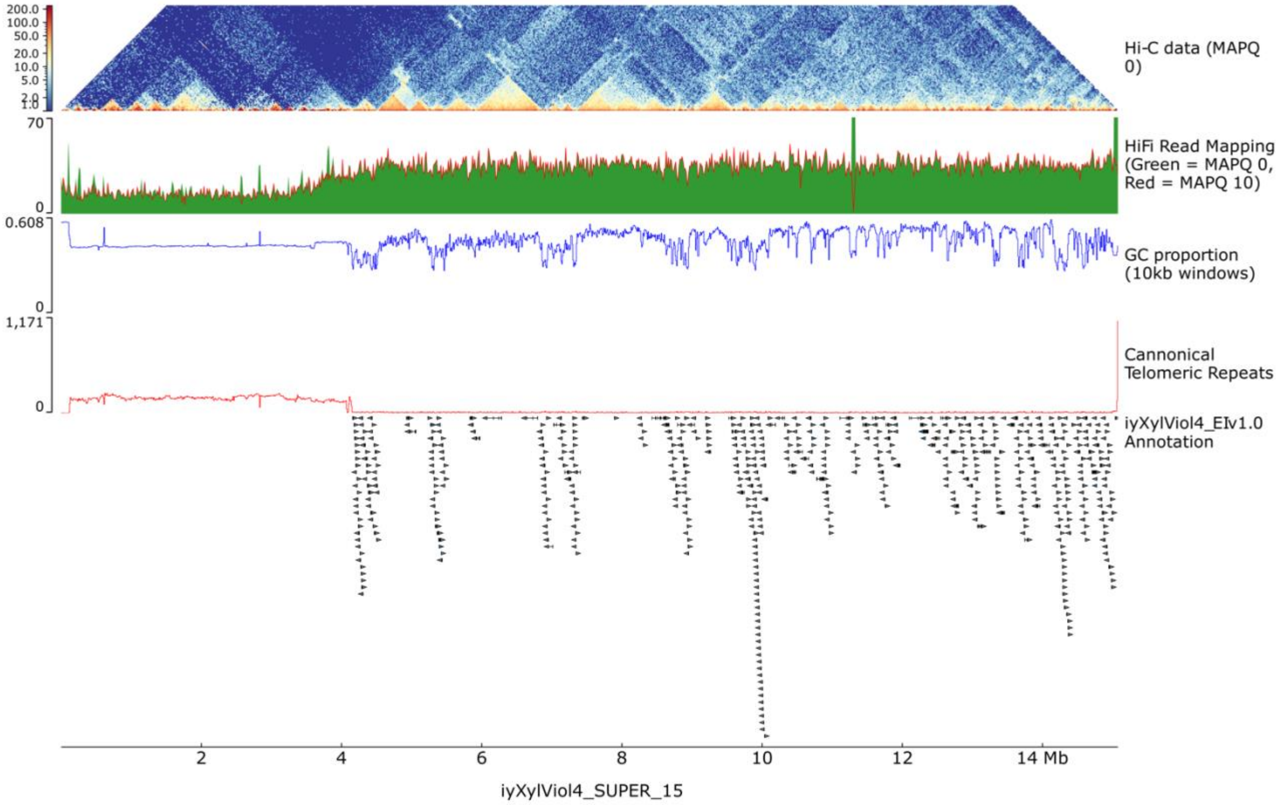
Hi-C data (MAPQ 0)
HiFi Read Mapping (Green = MAPQ 0, Red = MAPQ 10)
GC proportion (10kb windows)
Cannonical Telomeric Repeats
iyXylViol4_EIv1.0 Annotation
iyXylViol4_SUPER_15

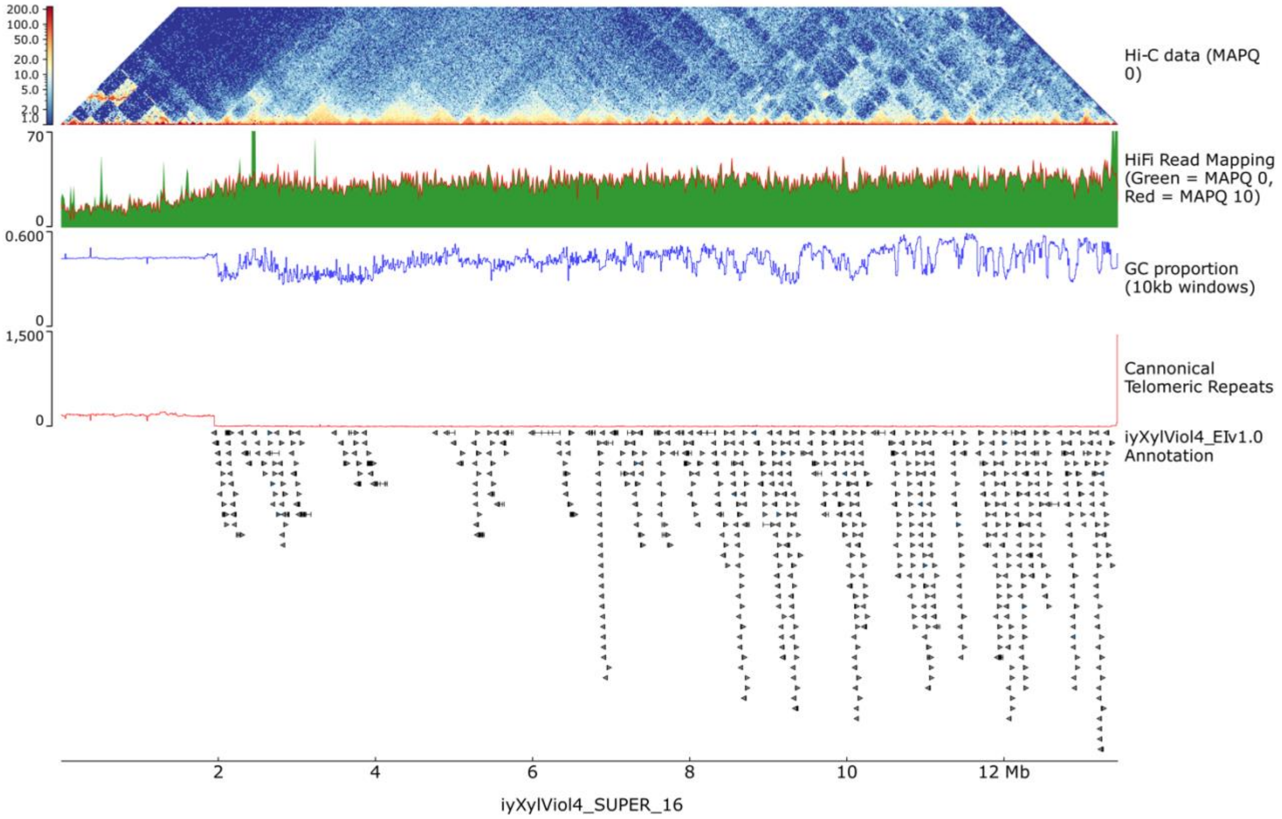
Hi-C data (MAPQ 0)
HiFi Read Mapping (Green = MAPQ 0, Red = MAPQ 10)
GC proportion (10kb windows)
Cannonical Telomeric Repeats
iyXylViol4_EIv1.0 Annotation
iyXylViol4_SUPER_16

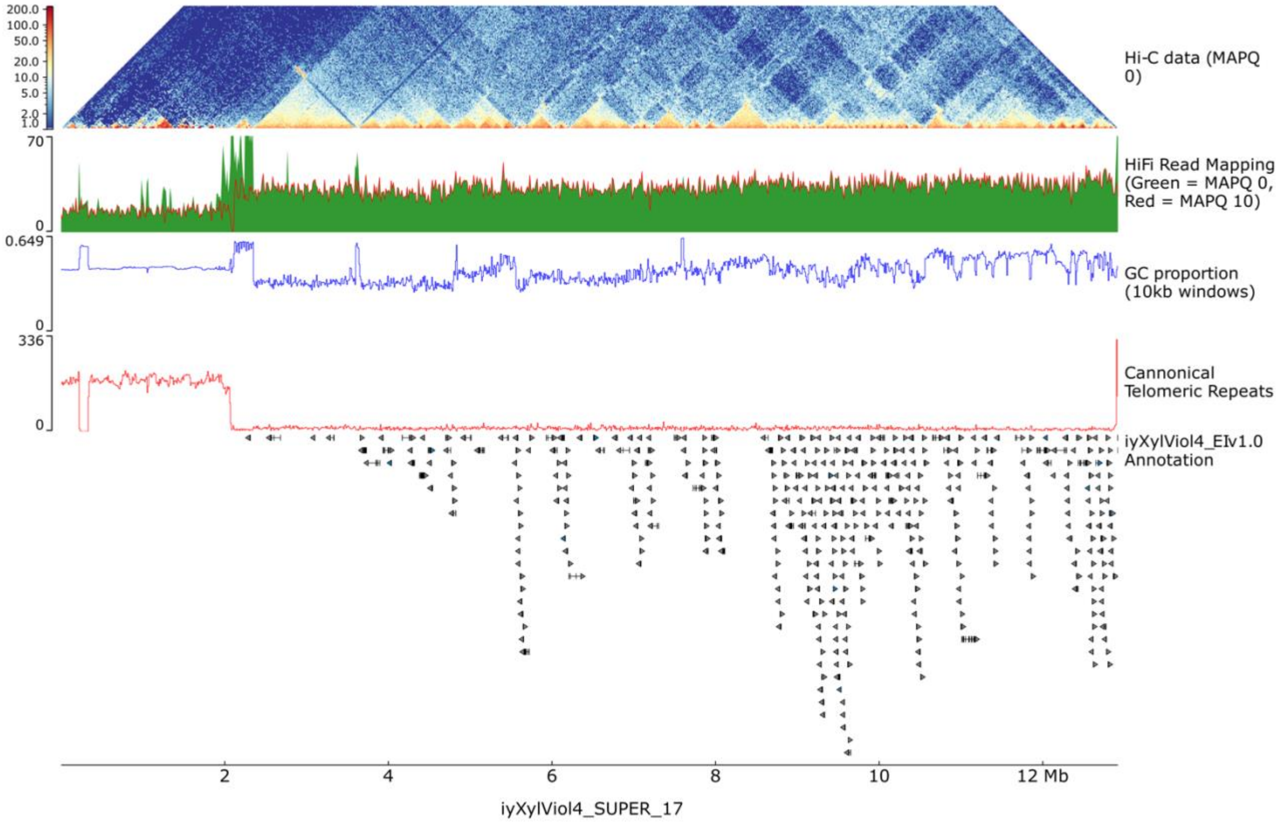
Hi-C data (MAPQ 0)
HiFi Read Mapping (Green = MAPQ 0, Red = MAPQ 10)
GC proportion (10kb windows)
Cannonical Telomeric Repeats
iyXylViol4_EIv1.0 Annotation
iyXylViol4_SUPER_17
2
4
6
8
10
12 Mb

#### Supplementary Figure 6. Visualisation of telomeric repeat diversity

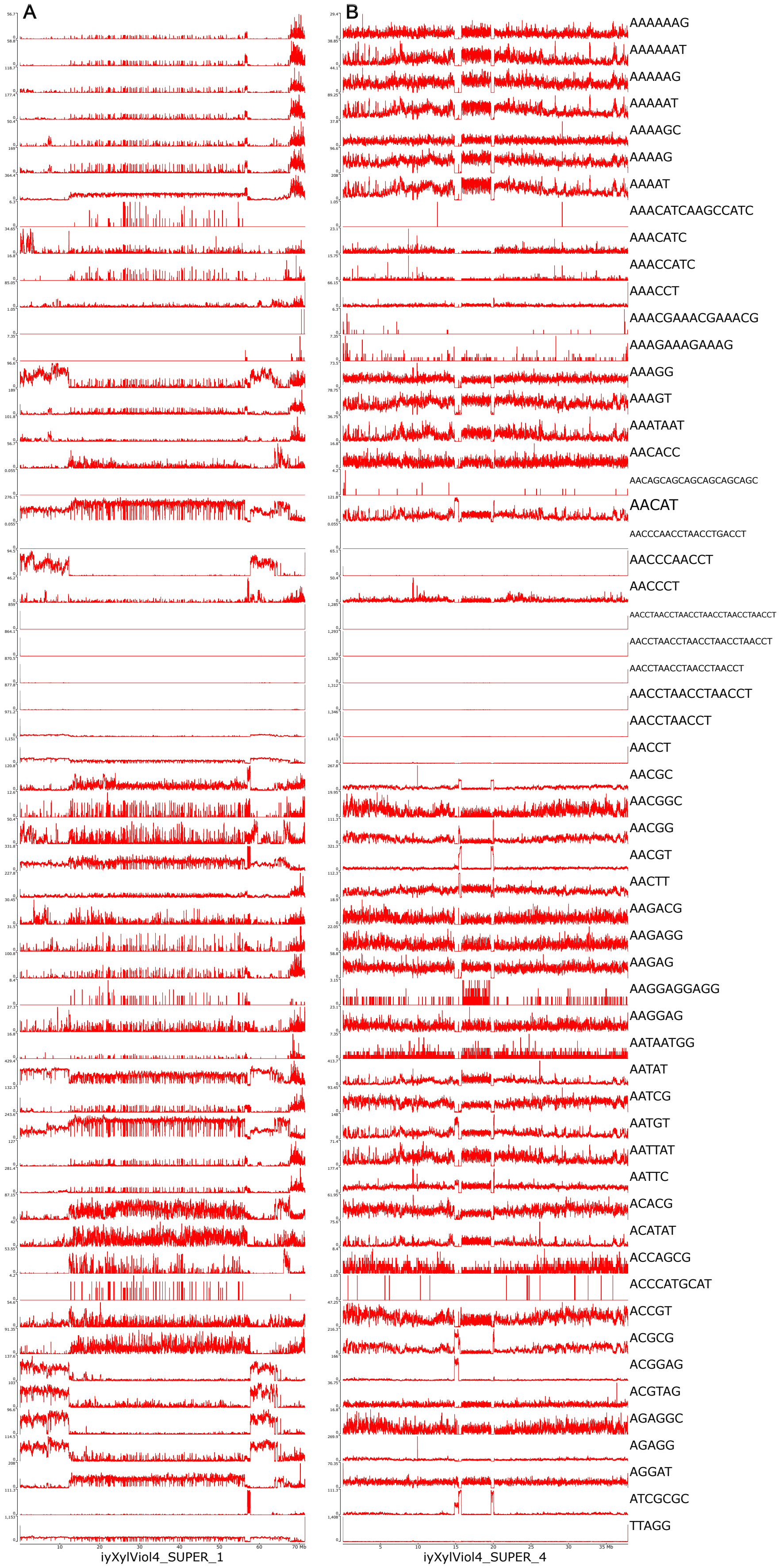


**Supplementary Figure 6. Telomeric Repeats in the iyXylViol4 genome assembly.** Exemplar plots of the distribution of repeat sequences identified using the `tidk explore` tool. **A)** iyXylViol4_SUPER_1, the largest pseudo-acrocentric pseudo-chromosomal unit, **B)** iyXylViol4_SUPER_4, the only fully resolved metacentric chromosome in the iyXylViol4 assembly.

#### Supplementary Figure 7. Percent identity, Satellite repeat, and monomer distribution on iyXylVio4_SUPER_1

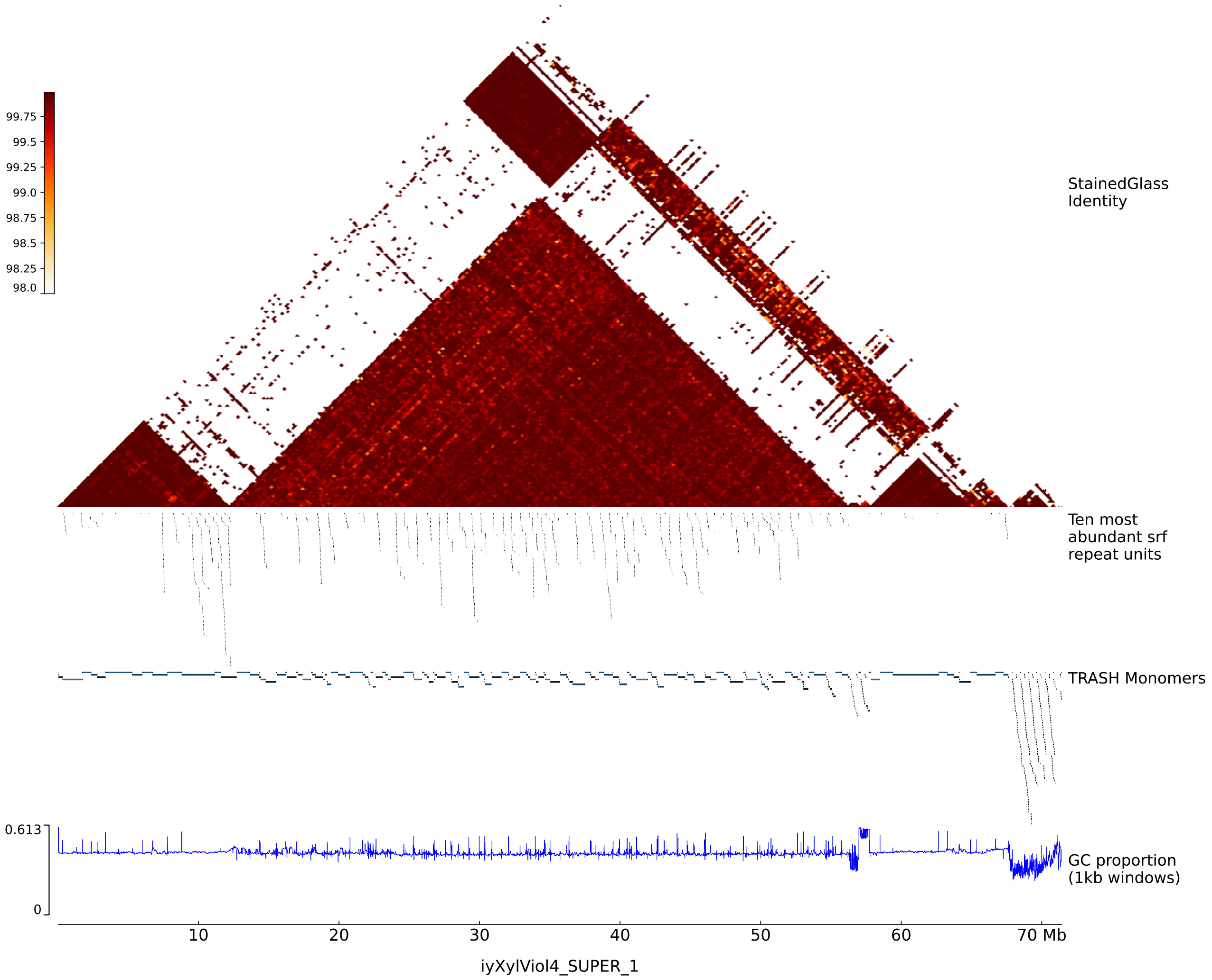


**Supplementary Figure 7. High identity repeat expansion on iyXylViol4 Super Scaffold 1. Top Panel:** Sequence identity, calculated using StainedGlass (Vollger et al. 2022), over the repeat expansion on iyXylViol4_SUPER_1. Range of Identities across the chromosome is 98-100%, with white being 0%. **Top middle panel:** Locations of ten most abundant (genome wide) Satellite repeat units identified using srf (Zhang et al. 2023). Details of the genome wide proportions of these units presented in Supplementary Table S6 **Bottom middle panel:** blocks monomeric repeat units identified using TRASH (Wlodzimierz, Hong, and Henderson 2023). Full results and coordinates of monomer mappings provided in Supplementary Table S7. **Bottom panel:** GC% proportion, plotted as above.

### Supplementary Figure 8. Repeat content of pseudo-chromosomal scaffolds

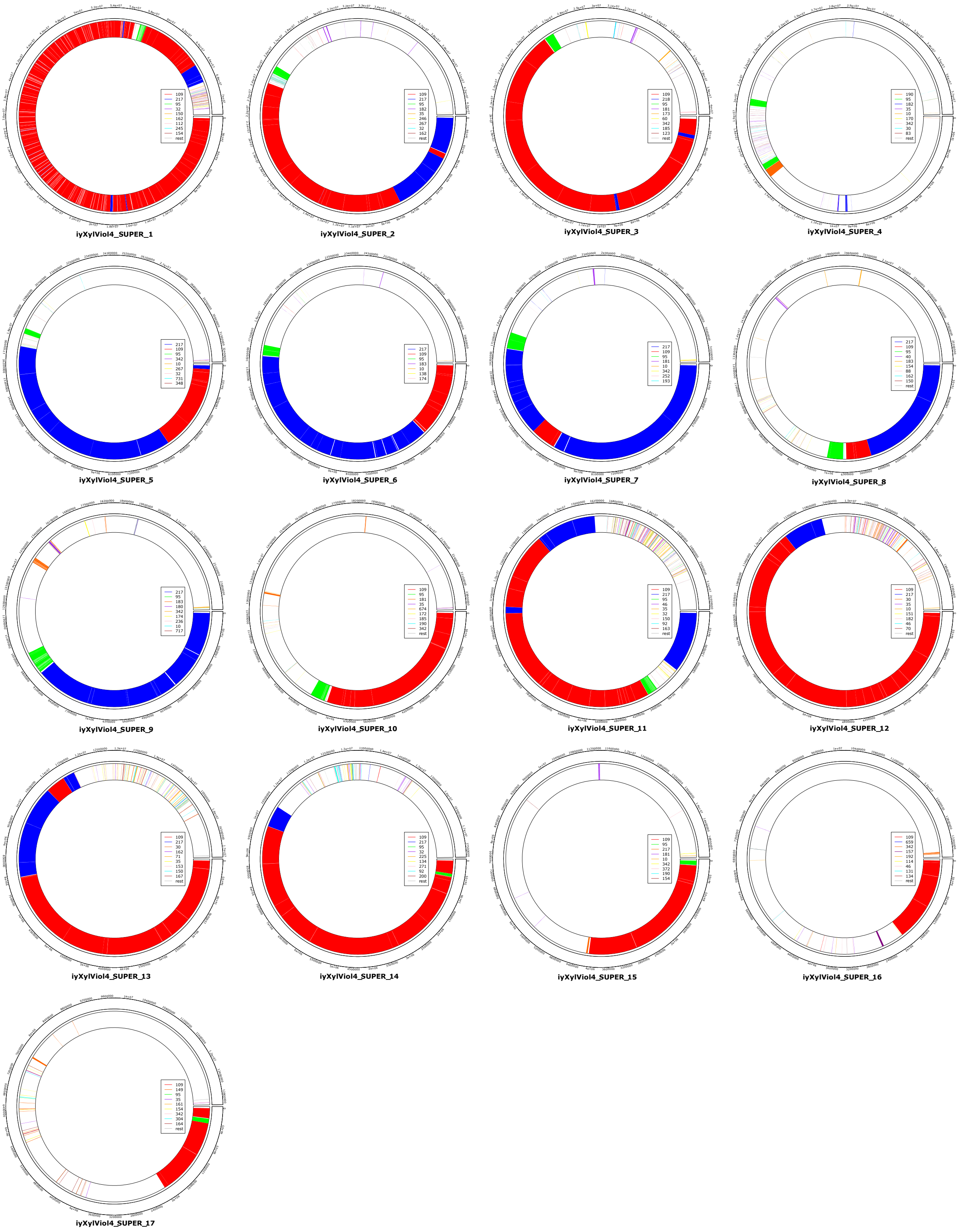


**Supplementary Figure 8. Repeat classification of pseudo-chromosomal scaffolds in the iyXylViol4 assembly.** Visualisation of the predominant satellite repeat monomers in the 17 pseudo-chromosomal super scaffolds of the iyXylViol4 assembly. The most abundant monomers are a 109mer (red), a 217mer (blue) or a 95mer (green). Other, less common, monomers are represented by different colours and differ between panels. Monomer identification and Circos plots conducted using TRASH (Wlodzimierz, Hong, and Henderson 2023), full results and coordinates of monomer mappings provided in Supplementary Table 7.
